# Spatial representation by ramping activity of neurons in the retrohippocampal cortex

**DOI:** 10.1101/2021.03.15.435518

**Authors:** Sarah A. Tennant, Harry Clark, Ian Hawes, Wing Kin Tam, Junji Hua, Wannan Yang, Klara Z. Gerlei, Emma R. Wood, Matthew F. Nolan

## Abstract

Neurons in the retrohippocampal cortices play crucial roles in spatial memory. Many retrohippocampal neurons have firing fields that are selectively active at specific locations, with memory for rewarded locations associated with reorganisation of these firing fields. Whether this is the sole strategy for representing spatial memories is unclear. Here, we demonstrate that during a spatial memory task retrohippocampal neurons encode location through ramping activity that extends within segments of a linear track approaching and following a reward, with the rewarded location represented by offsets or switches in the slope of the ramping activity. These ramping representations could be maintained independently of trial outcome and cues that mark the reward location, indicating that they result from recall of the track structure. During recordings in an open arena, neurons that generated ramping activity during the spatial memory task were more numerous than grid or border cells, with a majority showing spatial firing that did not meet criteria for classification as grid or border representations. Encoding of rewarded locations through offsets and switches in the slope of ramping activity also emerged in recurrent neural networks trained to solve a similar location memory task. Impaired performance of these networks following disruption of outputs from ramping neurons is consistent with this coding strategy supporting navigation to recalled locations of behavioural significance. We hypothesise that retrohippocampal ramping activity mediates readout of learned models for goal-directed navigation.

## Introduction

Spatial memory relies on complex brain network dynamics, with neurons in the hippocampus and associated retrohippocampal cortices (medial entorhinal cortex, pre- and para-subiculum) playing essential roles ^1^. Many neurons in these brain areas have receptive fields selective for location (Figure 1A). For example, place cells in the hippocampus and grid cells in retrohippocampal cortices have been widely studied ^1–3^, and are central to theories of navigation and memory (e.g. ^4–9^). However, theoretical studies suggest that continuous coding schemes in which neuronal firing rate is directly proportional to location (Figure 1B) may also be suited to control of navigational behaviours ^10–17^. While in principle such codes could be generated downstream from place or grid cells ^10–17^, it is unclear whether neurons in spatial memory circuits represent information using continuous codes or if coding schemes of this kind can be engaged by recall of memories.

**Figure 1.**
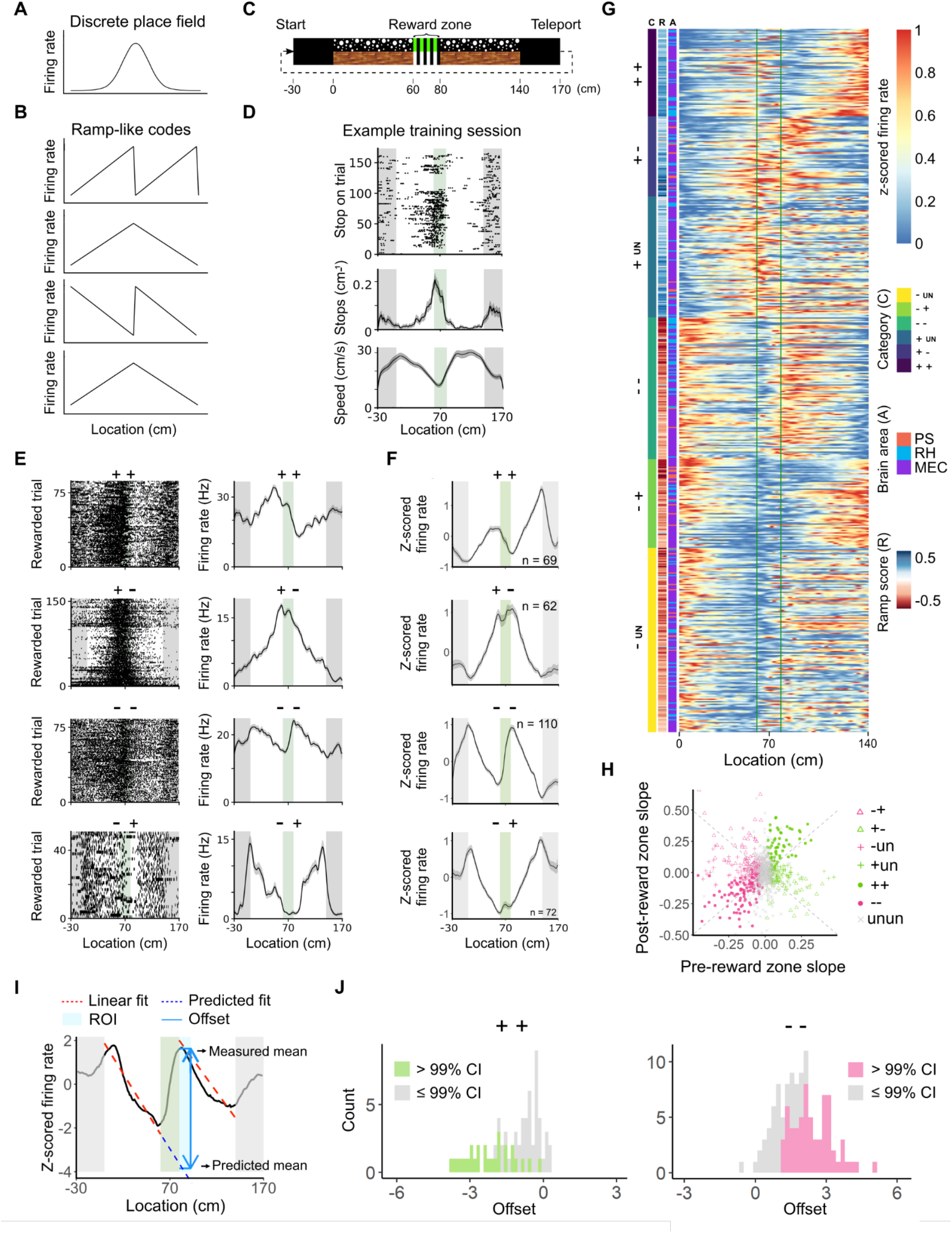
Interruption of ramping activity at a learned reward location. (A, B) Conceptualisation of discrete (A) and continuous (B) firing fields. (C) Schematic of the virtual track in the beaconed trial configuration. Numbers below the track indicate distance (cm) relative to the start of the track. (D) Example behaviour of a trained mouse on beaconed trials, showing a raster plot of stopping locations (upper), mean stops (per cm)(middle) and average running speed (lower) each as a function of position on the track (solid lines indicate mean values with standard error indicated in grey). The vertical green bar indicates the location of the reward zone. (E) Spike rasters (left) and the corresponding mean firing rate (right) as a function of track position for example neurons (solid lines indicate mean values with standard error indicated in grey). The first and third neurons are recorded from the training session shown in (D). Symbols above the plots indicate classification of the slope of the neuron’s ramping activity before and after the reward zone, e.g. + + indicates neurons with positive slope ramps on each side of the reward zone. (F) Z-scored firing rate as a function of position averaged across neurons classified by their ramp slope before and after the reward zone (symbols above each panel indicate classification). (G) Normalised firing rates as a function of location along the track (0 - 140 cm) for all neurons that have ramp-like firing in the region of the track before the reward zone. The start and end of the reward zone region is marked with vertical green lines. Neurons are categorised along the y axis by the direction of their slope before and after the reward zone (e.g. + +, - -, etc) and then ordered within each category according to their ramp slope before the reward zone. Columns to the left indicate the classification of activity based on fit of the linear model (C), the ramp score (R) and the location within the retrohippocampal region (L). (H) Slope of the fit to the firing rate on the track segment before the reward zone as a function of slope on the track segment after the reward zone. Dashed lines indicate points for which the slope would be identical on both track segments. Symbols indicate classification of individual neurons. (I) The ramp slope was estimated by fitting spiking activity as a function of location separately for the track segments before the reward zone (0 - 60 cm) and after the reward zone (80 - 140 cm). Offsets in firing rate were calculated by subtracting the actual firing rate in the first 5 cm after the reward zone with the firing rate predicted by extrapolating the fit from the track segment before the reward zone. (J) Distribution of offsets as calculated in (I) for neurons classified on beaconed trials as having ++ (left) or -- (right) activity patterns (++: t = -9.2, Df = 67, p = 1.9 × 10^-13^; --: t = 19.6, Df = 109, p < 2.2 x 10^-16^, t-test vs mean = zero). Coloured bars indicate individual neurons for which the offset was outside 99% confidence of intervals of the neuron’s predicted value.

Representations during learned spatial behaviours serve the purposes of tracking location along routes to a goal and of indicating whether the expected goal has been reached ^18^. Goal locations are associated with an increase in the density of hippocampal place fields ^19–23^ and with reorganisation of the firing fields of grid cells in the entorhinal cortex ^24,25^. Under these coding schemes, locations along the track and at goals are typically represented by different neurons, with the current location reflected in the identity of the currently active neurons. In contrast, under continuous coding schemes in which position is represented by firing rate, neural firing rates could ramp upwards or downwards during movement through an environment. If the initiation and termination of the ramp is then anchored to salient landmarks, which could include goal locations, then positions during movement between the anchoring landmarks may be represented by continuous changes in the firing rates of the ramping neurons. In this case, reaching a landmark could be indicated by cessation or reconfiguration of the ramping activity (Figure 1B). Coding schemes of this kind appear to be used by neurons in the striatum, which generate spatial ramping activity in anticipation of reward locations ^26–32^. Whether similar strategies are employed by neurons in the hippocampal formation is unclear.

To investigate coding schemes used by retrohippocampal neurons during recall of locations in learned environments, we used a memory task in which external sensory cues can be controlled on a trial by trial basis, enabling separation of cue driven and memory-dependent spatial representations ^33^. The task requires recall of the location of a reward zone on a virtual linear track (Figure 1C) ^33^. On cued trials the reward zone is marked by a prominent visual cue but otherwise the track lacks unique spatial cues. With training, mice efficiently obtain rewards by running directly to the reward zone (Figure 1D, Supplemental Figure 1) ^33^. This spatial behaviour is maintained when the cues marking the reward zone are removed, indicating that mice can solve the task by comparing online estimates of their distance from the start of the track, obtained using a path integration strategy, with estimates stored in memory ^33^. The path integration process may be driven either by sensory input or internal command signals. However, path integration alone is insufficient for accurate stopping unless the mouse also remembers where to stop. Thus, a memory for location on the track is critical to efficiently obtain rewards ^33^. For tasks with this structure, previously hypothesised continuous coding schemes ^10–17^ lead to predictions for neural representations that are distinct from and complementary to coding using classical discrete firing fields. First, under continuous coding schemes location may be represented by ramping neural activity. Second, the initiation and termination of ramping activity should be anchored to salient locations including the reward zone as well as the start and end of the track. Third, if ramping activity is driven by a neural path integrator then it should be maintained in the absence of location-specific cues. Fourth, if the locations that anchor the initiation or termination of ramping activity are recalled from memory, then transitions in firing at these locations should be maintained in the absence of cues that indicate the transition point.

We show here that in the retrohippocampal cortices of mice that have learned the location memory task, neurons encode location through ramping activity that either resets or switches polarity at the reward zone. Resetting and switches in ramp polarity were maintained after removal of reward zone cues, indicating that they can be driven by recall of the reward location. Neurons that generated ramping activity were observed more frequently than classical grid or border cells, and in open arenas a majority of these neurons had spatial firing fields that did not meet the criteria for classification as grid or border representations. Finally, we show that artificial recurrent neural networks can be trained to solve a similar location memory task and in doing so generate similar ramping representations that are interrupted at reward locations. Because successful activity patterns emerge in these artificial networks through reinforcement rather than by design, and as successful training did not require networks to be constrained by the specific connectivity of the retrohippocampal cortex, this suggests that ramping activity profiles that reset or change direction at salient landmarks may provide a general solution to the problem of representing remembered locations.

## Results

We used tetrodes targeted to retrohippocampal cortices, including the medial entorhinal cortex, presubiculum and parasubiculum (Supplemental Data 1), to record neural activity from mice carrying out the location memory task. Following spike sorting we obtained 1395 isolated single units (126.8 units / mouse, range 20-314; 11 mice) recorded from trained mice during sessions that contained a minimum of 30 correct trials (196.5 ± 7.7 trials / session; 187 sessions, 17 sessions / mouse, range 6-25). The tetrodes were usually lowered by ~ 50 - 100 μm following each session.

### Encoding of location by ramping activity

We focus our initial analyses on trials in which the animal stops in the reward zone, which we refer to as successful trials, and in which the reward zone was visible, which we refer to as beaconed trials. On these trials, a majority of recorded retrohippocampal neurons generated ramping activity within discrete regions bounded by the start of the track and the reward zone, and by the reward zone and the end of the track (n = 836 / 1395 neurons)(Figure 1E-G). For many neurons movement through the reward zone was associated with either a change in the slope of the ramp or resetting of the spike rate (Figure 1E-G). Changes in ramp slope were observed from positive before the reward to negative afterwards and vice versa (indicated as +- and -+ in Figure 1E-G). Resetting of the spike rate occurred as a step like reduction in spike rate of neurons with a positive ramp slope before and after the reward zone, and a step like increase for neurons with negative slopes before and after the reward zone (indicated as ++ and -- in Figure 1E-G).

To quantitatively evaluate ramp firing profiles, we classified neurons on the basis of their firing rate slopes in the track segments before and after the reward zone (Figure 1E-G). While there was a slight bias towards neurons with negative ramp slopes on the track segment before the reward zone, a continua of ramp slopes were observed and ramp slope on the track segment before the reward zone did not predict the ramp slope after the reward zone (Figure 1H). When neurons maintained a positive or negative ramp slope on both sides of the reward zone, their firing rate usually reset as the animal passed through the reward zone (Figure 1I-J). Classifications were similar when ramping activity was identified by fitting a linear model (p < 0.01 after correction for multiple comparisons, see Methods) or using a ‘ramp score’ based on correlations between position and firing rate (Figure 1G and Supplemental Figure 2A-C). Ramp-like activity was present in similar proportions of neurons from all animals (mean 60.7 ± 4.99 %, range 45.8 - 77.4), had firing rate peaks for ramps with positive slopes (or minima for ramps with negative slopes) that were strongly biased towards locations adjacent to the reward zone (Supplemental Figure 2D-E), was independent of theta modulation of spiking activity (Supplemental Figure 3), was similar in medial entorhinal cortex and pre/para-subiculum (Supplemental Figure 2F-G), and was present in only 2.02 % of shuffled data sets (n = 28235 / 1395000 shuffles)(Supplemental Figure 4). For ramping neurons, ramp slopes and track wide firing profiles were correlated between the first and second half of each recording session, indicating that ramp representations are stable at this time scale (ramp slope before the reward zone, r^2^ = 0.88, ramp slope after the reward zone, r^2^ = 0.77; track wide correlation, r^2^ = 0.47 ± 0.28). The corresponding correlation scores (r^2^ = 0.25, r^2^ = 0.21 and r^2^ = 0.23 ± 0.26), were substantially lower for non-ramping neurons (respectively: p < 2 x 10^-16^, F = 164.4, df = 1, ANOVA; p = 5 x 10^-15^, F = 63.0, df = 1, ANOVA; t = 14.6, df = 1103, p < 2 x 10^-16^, t-test).

To test whether ramping activity reflects position or other kinematic variables, we first fit firing rates in the track segment before the reward zone with generalised linear mixed effect models that included position, speed and acceleration as fixed effects (Figure 2A). For most neurons with ramping activity before the reward zone, their activity could best be accounted for by encoding of position, either alone or conjunctively with position as the dominant coefficient alongside speed and / or acceleration (n = 340/546 neurons with ramping activity before the reward zone)(Figure 2B-D). Ramping activity was also better accounted for by encoding of position rather than time (Supplemental Figure 5). As a further test, we evaluated neural firing rate profiles on trials in which mice failed to stop in the reward zone (Figure 3 and Supplemental Figure 6). We subdivided these miss trials into try trials, in which mice slowed down in the reward zone but didn’t stop, and run trials, in which mice maintained a high running speed within the reward zone (Figure 3A-B). Firing rate profiles of positionally modulated ramping neurons were similar for each trial outcome (Figure 3D, Supplemental Figure 6A), with firing rate slopes independent of trial outcome (Figure 3E) and offset behaviour of ++ and -- neurons maintained although somewhat reduced compared with hit trials (Figure 3F), indicating that for most neurons ramping activity, along with switches in slope and resetting of the firing rate in the region of the reward zone, is independent from changes in running speed associated with different trial outcomes.

**Figure 2.**
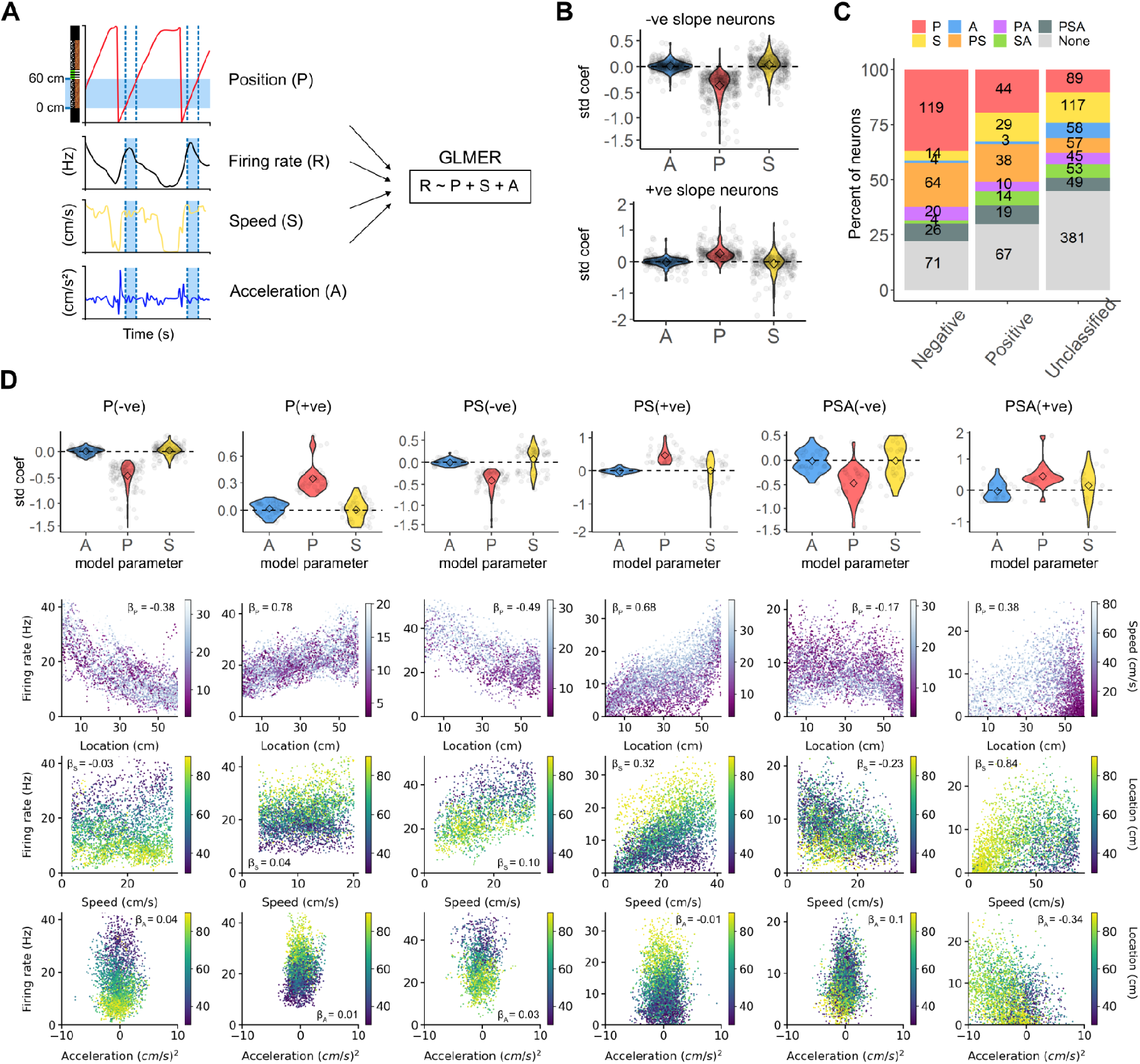
Differential influence of position, speed and acceleration on ramp-like firing. (A) Generalised linear mixed effect models that include speed (S), acceleration (A) and position (P) as fixed effects were fit to the firing rate in the track region approaching the reward zone (0-60 cm, see Figure 1C). The left panels show firing rate, position, speed and acceleration as a function of time for two consecutive trials from an exemplar experiment. The track region over which the model was fitted is indicated by the blue shaded bar. Speed and acceleration were calculated as the first and second derivatives with respect to time of the track position. Trial number was included as a random effect. (B) Standardised coefficients, which index the relative strength of each fixed effect variable, obtained from the model fits for neurons with negative (upper) and positive (lower) ramp slopes on the track segment before the reward zone. ANOVA indicates that the relative amplitude of coefficients for speed, acceleration and position differ (+ve slope: p < 2 x 10^-16^, Df = 2, 446, f = 93.18; -ve slope: p < 2 x 10^-16^, dF = 2, 642, f = 421.3; repeated measures ANOVA). For neurons in both groups the position coefficients were larger than those for acceleration (p < 2 x 10^-16^, Bonferroni corrected paired t-test) and speed (p < 2 x 10^-16^, Bonferroni corrected paired t-test). (C) Proportions of neurons for which each combination of P, A and S coefficients were significant at a threshold of p < 0.01 (Wald’s chi squared test) for neurons classified as having a positive slope, negative slope or as unclassified. (D) Distribution of standardised coefficients (upper row) and example data (lower rows) for neurons with positive (+ve) or negative (-ve) rap slopes and classified as modulated solely by position (P), by position conjunctively with speed (PS), and position conjunctively with speed and acceleration (PSA). Example data shows firing rate as a function of position (second row), running speed (third row) and acceleration (lower row). Firing rates as a function of position are colour coded by speed (cm/s) while firing rates as a function of speed or acceleration are colour coded by position along the track (0 - 60 cm).

**Figure 3.**
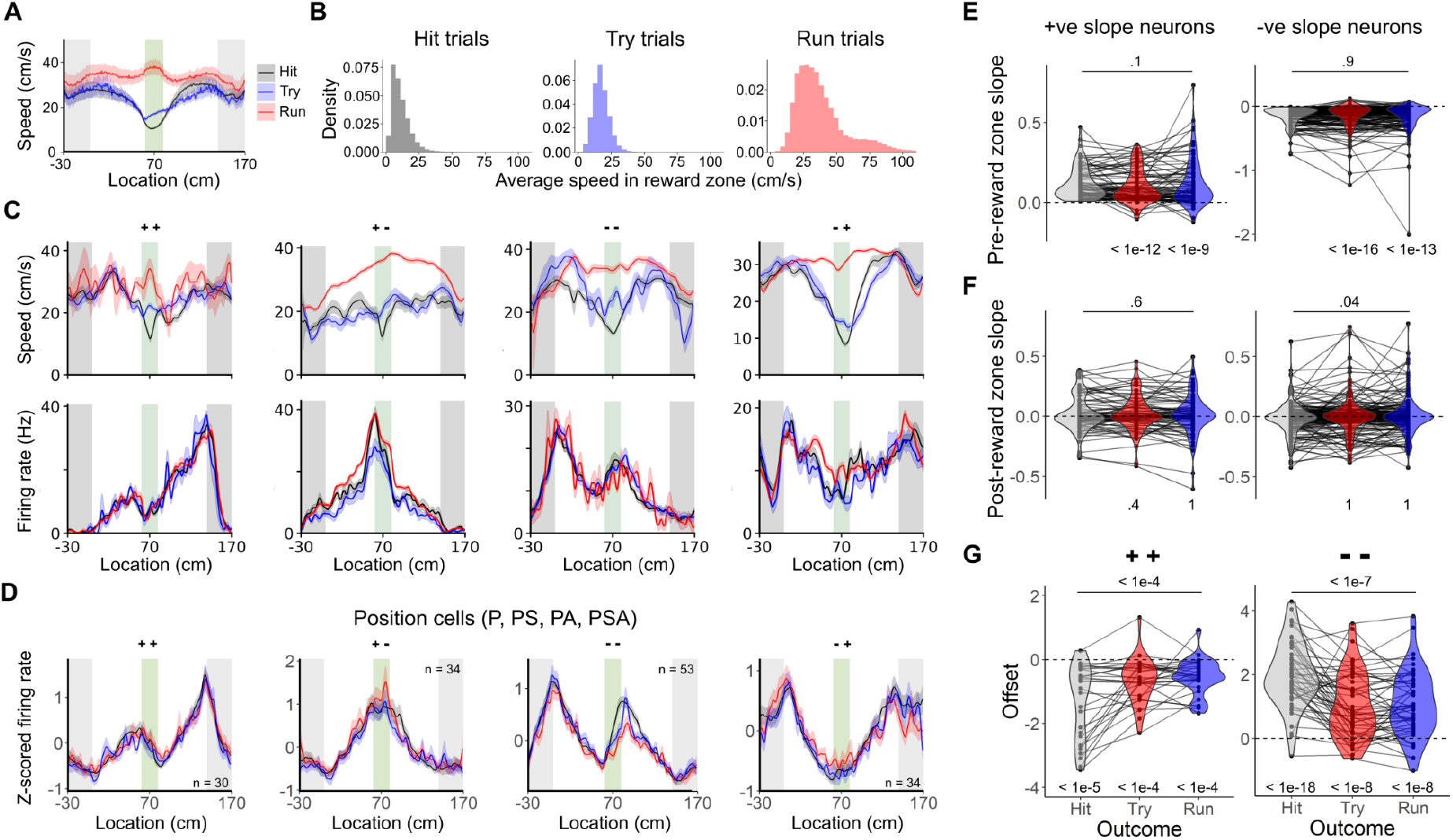
Relationship between task performance and ramp-like activity. (A) Population average of mean running speed as a function of position for each trial outcome across all animals (n = 11). Trials were classified as ‘hit trials’ when mice stopped in the reward zone region, as ‘try trials’ when mice slowed down to < 95th percentile of reward zone speeds in hit trials but did not stop, and as ‘run trials’ when mice maintained a speed > 95th percentile of reward zone speeds in hit trials. Shaded regions correspond to the standard error of the mean across animals. Note that given the reversal of the speed profile between hit and try groups (reduced in the reward zone) compared with the run group (higher in the reward zone), neurons with activity determined exclusively by running speed should show a similar reversal. (B) Trial level distribution of mean running speeds within the reward zone for hit, try and run trial outcomes. The histograms are calculated from all sessions (n = 187) and all trials with a given outcome (hit: n = 12130; try: n = 1534; run: n = 1111). (C) Examples of running speed (upper) and firing rate (lower) as a function of track position. Note that for some animals the average running speed in the hit group can be greater than zero because the stopping location varies between trials. Shaded regions correspond to the standard error of the mean across trials. (D) Population averaged firing rates as a function of position on hit, try and run through trials for positionally modulated neurons (P, PS, PA, PSA groups). The neurons were classified on hit trials as having +-, -+, ++ and -- firing rate profiles. Shaded regions correspond to the standard error of the mean across cells. (E) Slope of positive and negative ramping activity on the track segment before the reward zone as a function of trial outcome. There was no significant effect of trial outcome (+ve slopes: Df = 2, 164, F = 2.1, p = 0.12; -ve slopes: Df = 2, 308, F = 0.069, p = 0.93; repeated measures ANOVA) and for all outcomes the slope values differed significantly from zero (p < 2 x 10^-10^, Bonferroni corrected t-test). (F) Slope of positive and negative ramping activity on the track segment after the reward zone as a function of trial outcome. There was no significant effect of trial outcome (+ve slopes: Df = 2, 308, F = 3.2, p = 0.04; post-reward zone -ve slopes: Df = 2, 164, F = 0.427, p = 0.65; repeated measures ANOVA). (G) Offset between the predicted and actual firing rate following the reward zone as a function of trial outcome for neurons with positive slopes before and after the reward zone (++) and for neurons with negative slopes before and after the reward zone (--). The extent of the offsets varied according to trial outcome (++: Df = 2, 56, F = 12.43, p = 3.4 x 10^-5^; --: Df = 2, 102, F = 21.91, 1.2 x 10^-8^), but for all outcomes the offsets differed from zero (p < 7 x 10^-5^, Bonferroni corrected t-test). Differences between offsets may reflect increased trial to trial variation in the location at which resetting occurs in some neurons (Supplemental Figure 6).

Together, these data indicate that firing rate profiles in which ramping activity encodes location within discrete track segments and reconfigures at a reward location are a common feature of neural activity in retrohippocampal cortices. These activity patterns contrast with discrete firing fields that are typical of place and grid cell fields recorded previously on linear virtual tracks ^34–36^.

### Ramp-like firing rate profiles in the absence of reward zone cues

To test whether changes in the slope or resetting of the rate of ramp firing at the reward zone results from recall of a memory of the track structure, we investigated activity on probe trials in which rewards and the visible cues indicating the reward zone were omitted (1/10 trials were probe trials, interleaved with 4/5 trials that had the standard cue configuration and 1/10 trials that are rewarded but lack the visible cue)(Figure 4A-B). We focus on neurons with firing rate profiles that depend on track position, either alone or conjunctively with speed or acceleration, and for which there were sufficient numbers of probe trials (see Methods) (n = 97 / 240 neurons with positional ramping before the reward zone, 38 sessions, 9 mice).

**Figure 4.**
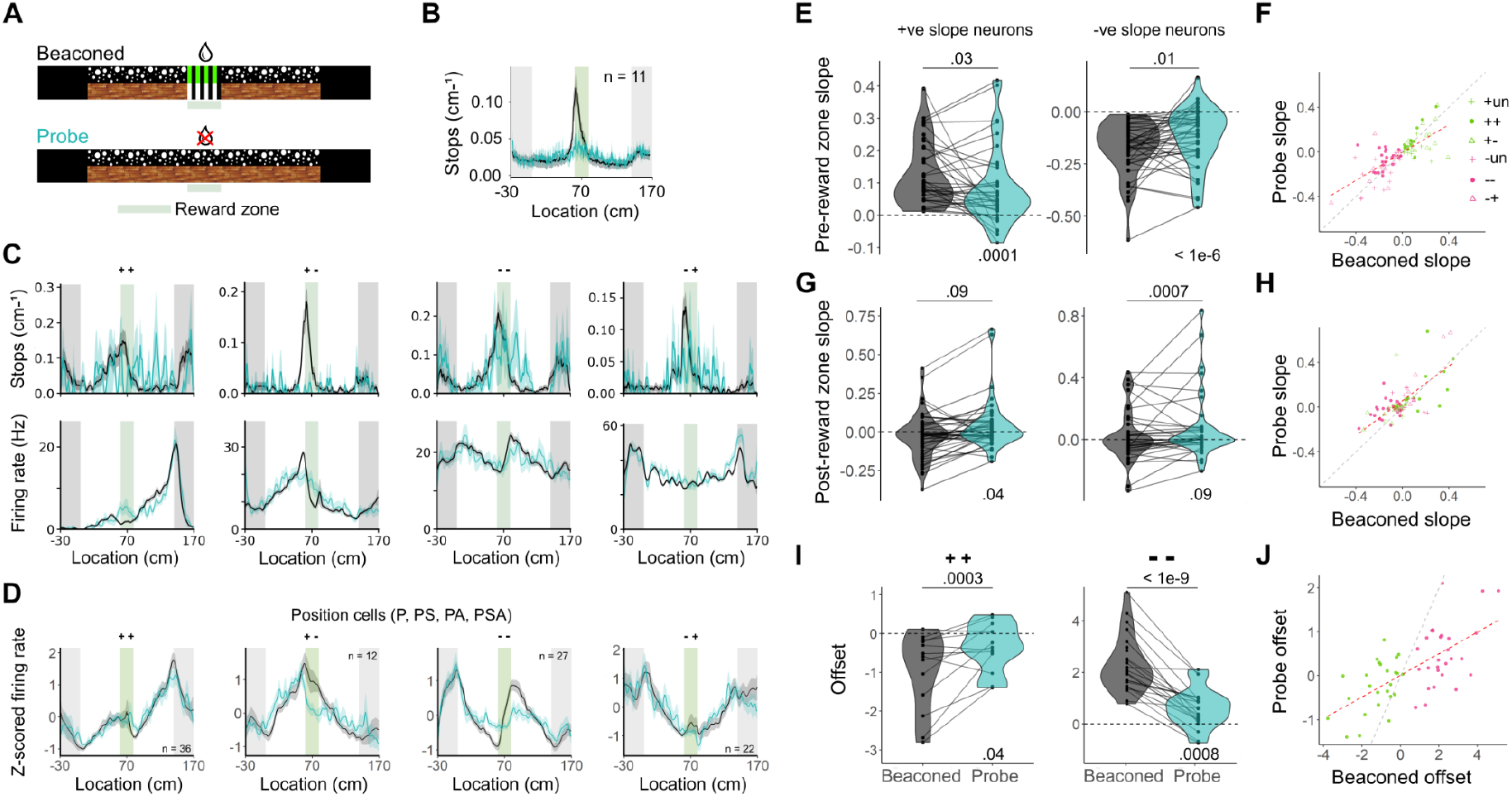
Location cues are not required for reconfiguration of ramping firing rate profiles at the reward zone. (A) Schematic of track configuration. (B) Population mean of behaviour from trained mice on beaconed and probe trials, showing the mean stops (per cm) as a function of position on the track. Shaded regions correspond to the standard error of the mean across animals. (C) Examples from beaconed trials (black) and probe trials (blue) of stop histogram (upper row) and mean firing rate (lower row, shaded areas indicate the SEM) as a function of track position. The neurons were classified (++, +-, --, -+) according to their firing rate profiles on beaconed trials. (D) Mean firing rates as a function of track position on beaconed trials (black) and probe trials (blue) for the populations of neurons classified with ++, +-, --, -+ firing rate profiles. (E, G) Slopes of ramping activity on the track segment before the reward zone (E) and after the reward zone (G) differed between beaconed and probe trials (pre-reward zone +ve slopes: t = 2.3, df = 36, p-value = 0.028; pre-reward zone -ve slopes: t = -2.6, df = 45, p-value = 0.012; post-reward zone +ve slopes: t = -1.7, df = 36, p-value = 0.093; post-reward zone -ve slopes: t = -3.7, df = 45, p-value = 0.00067; paired Student’s t-tests). Slopes before the reward zone on probe trials nevertheless remained above / below zero on for +ve / -ve slope neurons respectively (+ve: t = 4.3, df = 36, p-value = 0.00011; -ve: t = 2.1, df = 36, p-value = 0.038; one sample t-test versus mean of zero). Lines connect points from the same neuron. (F, H) On track segments before (F) and after (H) the reward zone, ramp slopes on probe trials varied as a function of the beaconed trial slope (pre-reward zone: adjusted R^2^ = 0.50, F = 83.9 on 1 and 81 DF, p = 3.9 × 10^-14^; post-reward zone: adjusted R^2^ = 0.45, F = 67.3 on 1 and 81 DF, p = 2.9 × 10^-12^). (I) Firing rate offsets for ++ and -- neurons, measured as in Figure 1H, were reduced on probe compared to beaconed trials (++: t = -4.3, df = 23, p-value = 0.00027; --: t = 9.1, df = 30, p-value =s; paired t-test), but remained greater than expected from linear extrapolation of ramping activity before the reward zone (++: t = -2.19, df = 23, p-value = 0.039; --: t = 3.7, df = 30, p-value = 0.00078; one sample t-test). (J) Firing rate offsets on probe trials were correlated with offsets on beaconed trials (adjusted R^2^ = 0.3938, F = 52.97 on 1 and 79 DF, p = 2.2 x10^-10^).

The effects of cue removal differed between positional ramping neurons. For many neurons, encoding of the reward zone, by changes in the slope of ramping activity or by resetting of the spike rate, was maintained on probe trials (Figure 4C, Supplemental Figure 6B). This stability was manifest in similarity between average firing rate profiles obtained on beaconed and probe trials from populations of positional neurons grouped according to their slopes before and after the reward zone (e.g. ++, +-, -+ and --)(Figure 4D). To assess the effects of cue removal across the neuronal population we determined the effects on ramp slopes and offsets. Ramp slopes showed variable responses to removal of the cues, with some neurons maintaining a similar slope, which is consistent with their activity being driven by a path integration mechanism, and others showing a reduced slope, which is consistent with the reward zone cues driving their activity (Figure 4E, G). The latter group of neurons drove a reduction in the mean ramp slope on probe compared to beaconed trials, although the population slopes remained substantially above chance (Figure 4E, G) and slopes on probe trials were correlated with slopes on beaconed trials (pre-reward zone: adjusted R^2^ = 0.50, p = 3.9 × 10^-14^; post-reward zone: adjusted R^2^ = 0.45, p = 2.9 × 10^-12^)(Figure 4F, H), which is consistent with the maintained ramping profiles in the average activity. The effects of cue removal on the offset in ramping activity was also consistent with variation between neurons in the influence of path integration and cue-driven mechanisms. Thus, offsets in ramping activity of ++ and -- neurons were maintained on probe trials, although were reduced compared to probe trials (Figure 4I), but again were strongly correlated between the two trial types (adjusted R^2^ = 0.39, p = 2.2 x 10^-10^)(Figure 4J).

The maintenance of ramping profiles in the absence of reward zone cues indicates that recall of the track structure is sufficient for ramping activity and its interruption at the reward zone. Reduced ramping and offset scores of some neurons suggests that cues associated with the reward zone also play a role in establishing firing rate profiles. These observations are consistent with activity of many neurons in the MEC reflecting readout of an internally generated learned model of the track, with other neurons being driven directly by visual cues indicating the reward zone.

### Ramping neurons have spatial firing fields in open environments

Retrohippocampal cortices are widely associated with grid, border and head direction representations ^37–41^. However, because ramp firing patterns that we report here are recorded during a task in which recall of the track locations is necessary for reward, whereas firing properties of retrohippocampal neurons are usually evaluated in the absence of location-specific reward contingencies, it’s unclear which of the previously described functional cell types ramping neurons correspond to. To address this, in a subset of behavioural sessions after recording from neurons in the location memory task we recorded from the same neurons in an open arena (Figure 5A). We used a hierarchical scheme to classify cells ^37^, with rank grid > border > other spatial > head direction > non-spatial. In this scheme, cells were assigned to the highest ranked category for which their identity score was greater than the 99^th^ percentile of their corresponding shuffled data (see Methods). For most neurons with positional ramping activity in the location memory task, their firing in the open arena was classified in the other spatial category (n = 145 / 201)(Figure 5B-D). Positional ramping neurons that had border, grid or pure head direction firing patterns in the open field were found relatively infrequently (n = 10, 7 and 18 / 201 respectively). The proportion of border, grid and pure head direction cells was similar among ramping and non-ramping neurons (Figure 5C, Supplemental Figure 7A-B), and the proportion of ramping and non-ramping neurons was similar among each open arena cell type (Figure 5D). The distribution of speed modulated activity in the open arena was also similar among ramping and non-ramping neurons (Supplemental Figure 7B).

**Figure 5.**
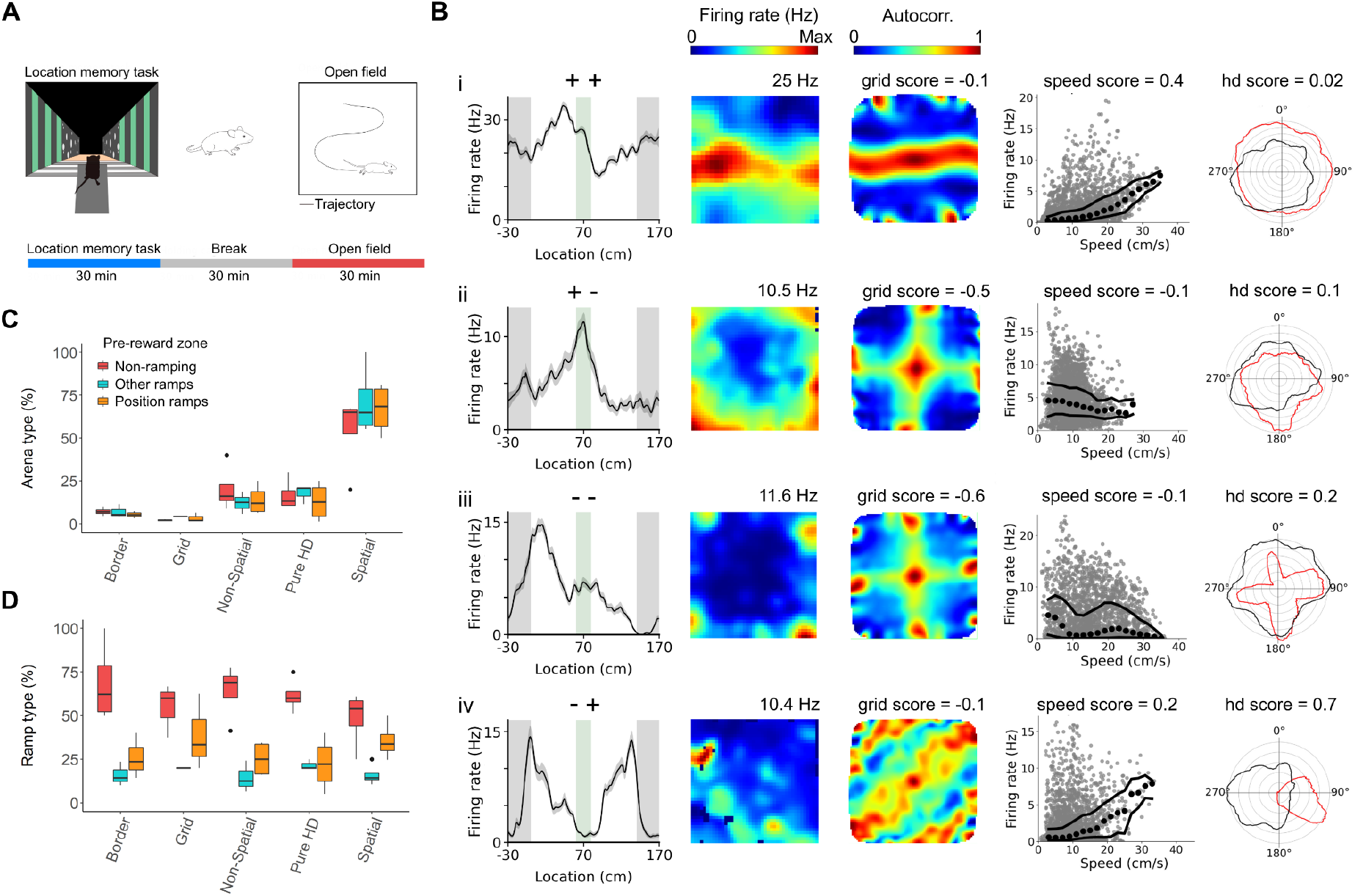
Ramping neurons have diverse spatial firing patterns in an open arena. (A) Animals performed the location estimation task for ~30 minutes, were placed in a holding arena for ~30 minutes, and then explored the open arena for ~30 minutes. (B) Example activity in the open arena of cells with ramping activity on the linear track characterised by +ve slopes approaching and following the reward zone (i), +ve slope approaching and -ve slope following the reward zone (ii), -ve slopes approaching and following the reward zone (iii) and -ve slope approaching and +ve slope following the reward zone (iv). Activity in the open arena was classified as either spatial (i, iv) or border (ii, iii). Panels show (from left to right): mean firing rate on the virtual track as a function of position; spatial heat maps of firing rates in the open arena; spatial autocorrelograms of the open arena firing rate; firing rate in the open arena as a function of speed; and polar plots of average firing rate in the open arena as a function of head direction (red) and movement (black). (C) Proportion of each open arena cell type (border, grid, head direction, non-spatial, other spatial) among neurons with activity on the linear virtual track classified as position-dependent ramping, other ramping, and non-ramping. (D) Proportion of neurons classified on the linear virtual track classified as position-dependent ramping, other ramping, and non-ramping among neurons classified as border, grid, head direction, non-spatial, other spatial. In (C) and (D), boxes show the median, 25th and 75th percentiles, and whiskers extend to the largest value 1.5 times the interquartile range from the box. Points outside this range are plotted individually.

We also evaluated the correspondence between the activity of ramping neurons and features used to distinguish principal cells from interneurons. The majority of ramping neurons had firing properties characteristic of principal cells, while a small proportion had firing properties that are typical of interneurons (Supplemental Figure 7C). Ramp firing of putative interneurons usually had a negative slope on the track segment before the reward zone, with few putative interneurons having positive ramp slopes (Supplemental Figure 7D).

Together, these data indicate that the population of neurons with ramping activity is largely, but not exclusively, non-overlapping with grid, head direction and border cell populations, while many neurons that in an open arena lack well defined spatial firing fields may participate in representation of learned information by generating ramp-like representations that interrupt around recalled reward locations.

### Recurrent neural networks learn to estimate location using ramp-like representations

If initiation and termination of ramping representations at behaviourally important locations is an effective representation for guiding learned behaviours, then similar activity profiles may emerge in artificial neural networks trained to solve location memory tasks. To investigate this, we trained a network model that received visual input corresponding to a virtual track with layout similar to the experimental track. To promote cue-independent strategies we trained the network solely with tracks in which the visual reward zone cues were absent. The model network generated stop or go signals to control movement along the track. The model was organised such that visual signals were processed by a three layer convolutional neural network with the output fed into a recurrent neural network that generated movement control signals (Figure 6A). We chose this network architecture because recurrent connections are a feature in many handcrafted models of the entorhinal cortex ^42^ and we focussed on visual inputs as visual cues delineate the segments of the experimental track and visual signals are encoded by neurons within the entorhinal cortex ^43^. Networks trained with a reinforcement learning algorithm ^44^ successfully learned the task (Figure 6B). Before training, the agent controlled by the network stopped frequently on the track with no particular spatial stopping pattern (Figure 6C). After training, the agent stopped at the reward zone on non-beaconed (Figure 6D) and probe trials (Figure 6E) indicating that the network solves the task using a distance-based strategy. The agent’s stopping patterns were remarkably similar to those of mice before and after learning the task ^33^.

**Figure 6.**
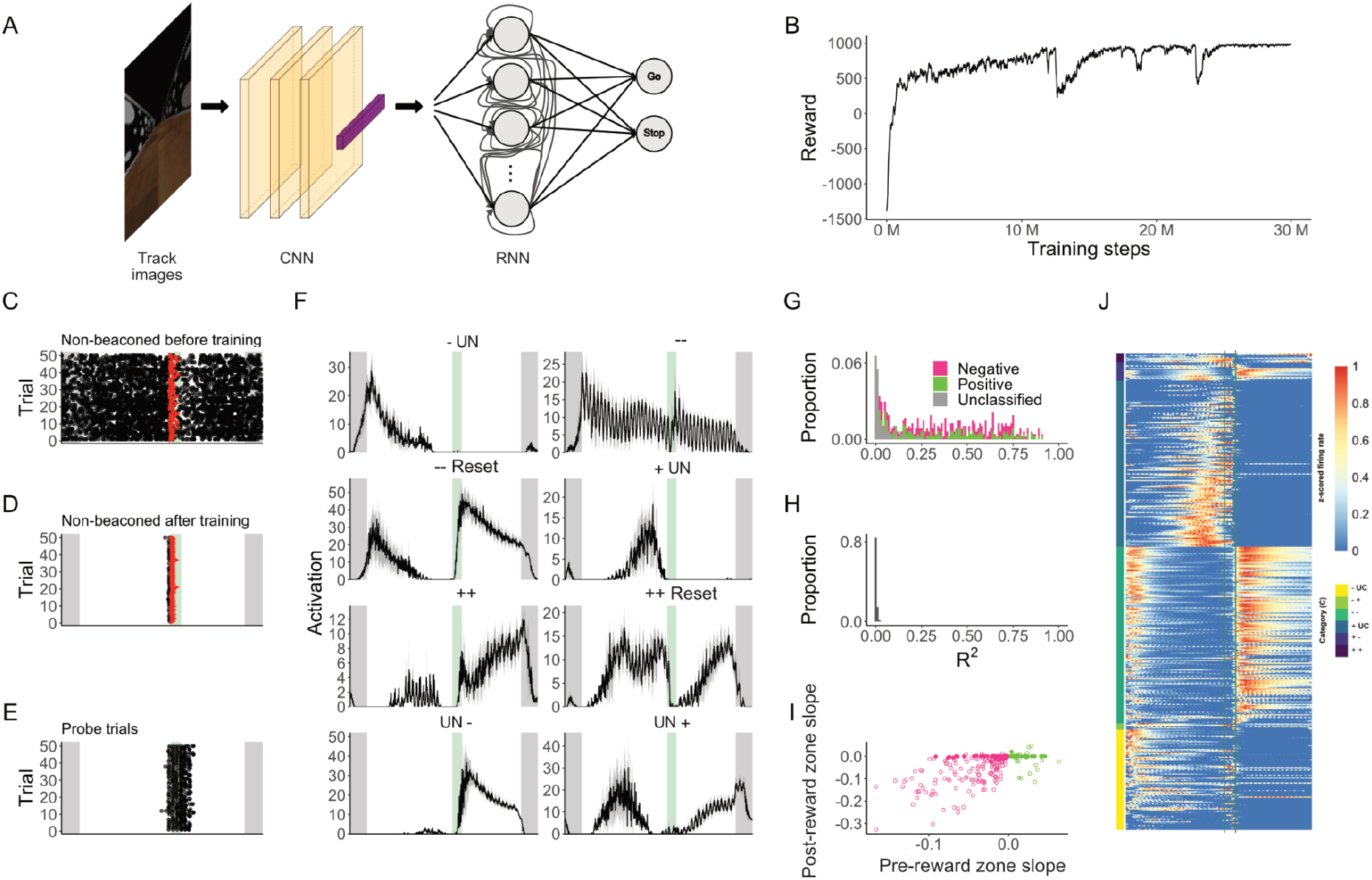
Recurrent networks trained to estimate location generate ramp-like representations. (A) Schematic of the network. Images of the track were played into a convolutional neural network (CNN) that sends outputs to a recurrently connected network (RNN) that sets the activity of movement control neurons. On each simulation step the network can either select GO, which advances the agent, or STOP, which causes the agent to remain at the same location. During training a reinforcement learning algorithm modifies the network weights according to whether the agent stops at the reward zone. (B) Reward obtained as a function of training step. If the agent stops in a non-goal location, it receives a reward of -1, whereas moving forwards is a neutral action with a reward of 0, and stopping in the goal locations yields a reward of 1000. Dips in the reward occur when the agent tests new (unsuccessful) navigation strategies. After developing a successful strategy during training, the agent typically takes ~ 250 simulation steps to reach the end of the track. (C-D) Rasters of stopping locations on non-beaconed trials before (C) and after (D) the agent has learned the task. (E) Rasters of stopping locations on probe trials after the agent has learned the task. (F) Examples of RNN unit activity as a function of track location after the agent has learned the task. (G) R^2^ values for fits of activity of RNN units as a function of position in the track segment before the reward zone. Units are classified as in Figure 1. (H) As for (G) but for shuffled data. (I) Slopes of the firing rate of RNN units after the reward zone as a function of slopes before the reward zone. On the track segment before the reward zone 43.4% of neurons were classified as having a negative ramp slope and 29.3% a positive ramp slope. On the track segment after the reward zone 33.0% were classified as having a negative slope and 4.7% a positive slope. (J) Normalised firing rate as a function of position on the track for all units in the RNN.

As in experimental recordings from retrohippocampal neurons, units in the trained recurrent network showed ramping activity that was interrupted at the reward zone (Figure 6F, J). The ramping activity of units in the recurrent network was clearly distinguishable from activity patterns generated by shuffled datasets (Figure 6G,H), and had either a positive or a negative slope (Figure 6F, I-J), with the slope on the region of the track approaching the reward zone not predicting the slope after the reward zone (Figure 6I). Also similar to the experimental data, ramps generated by the trained recurrent network more frequently had a negative rather than a positive slope, although in the recurrent networks negative slopes were typically steeper than positive slopes (Figure 6I).

The presence in artificial recurrent neural networks and in the retrohippocampal cortices of activity profiles characterised by ramping activity and its interruption at reward locations is consistent with these activity patterns providing an effective readout of location. To test whether ramping activity of neurons in the trained networks contributes to successful completion of the task, we explored the consequences of perturbing neurons in the recurrent network targeted according to their activity pattern (Figure 7A). We first tested blocking of the output from either ramping or non-ramping neurons to other neurons in the recurrent network. In contrast to the trained network, which stops at the reward location on all trials (Figure 7B), networks in which recurrent outputs from either ramping neurons (Figure 7C) or non-ramping neurons (Figure 7D) were inactivated did not show a spatial stopping strategy, indicating that activity of both neuron types is critical for the network to function. Because inactivation might be expected to cause substantial behavioural deficits simply through disrupting recurrent network activity, we evaluated two further manipulations. Selective block of outputs from ramping neurons to downstream movement control neurons reduced the spatial selectivity of the stopping behaviour (Figure 7E), whereas the spatial selectivity was largely maintained when the corresponding outputs from non-ramping neurons were blocked (Figure 7F). Clamping the activity of either ramping or non-ramping neurons to their mean activity level across a trial also abolished spatial stopping behaviour, resulting in either abolished stopping when the activity of ramping neurons was clamped (Figure 7G), or a greater frequency and modified distribution of stops when activity of non-ramping neurons was clamped (Figure 7H).

**Figure 7.**
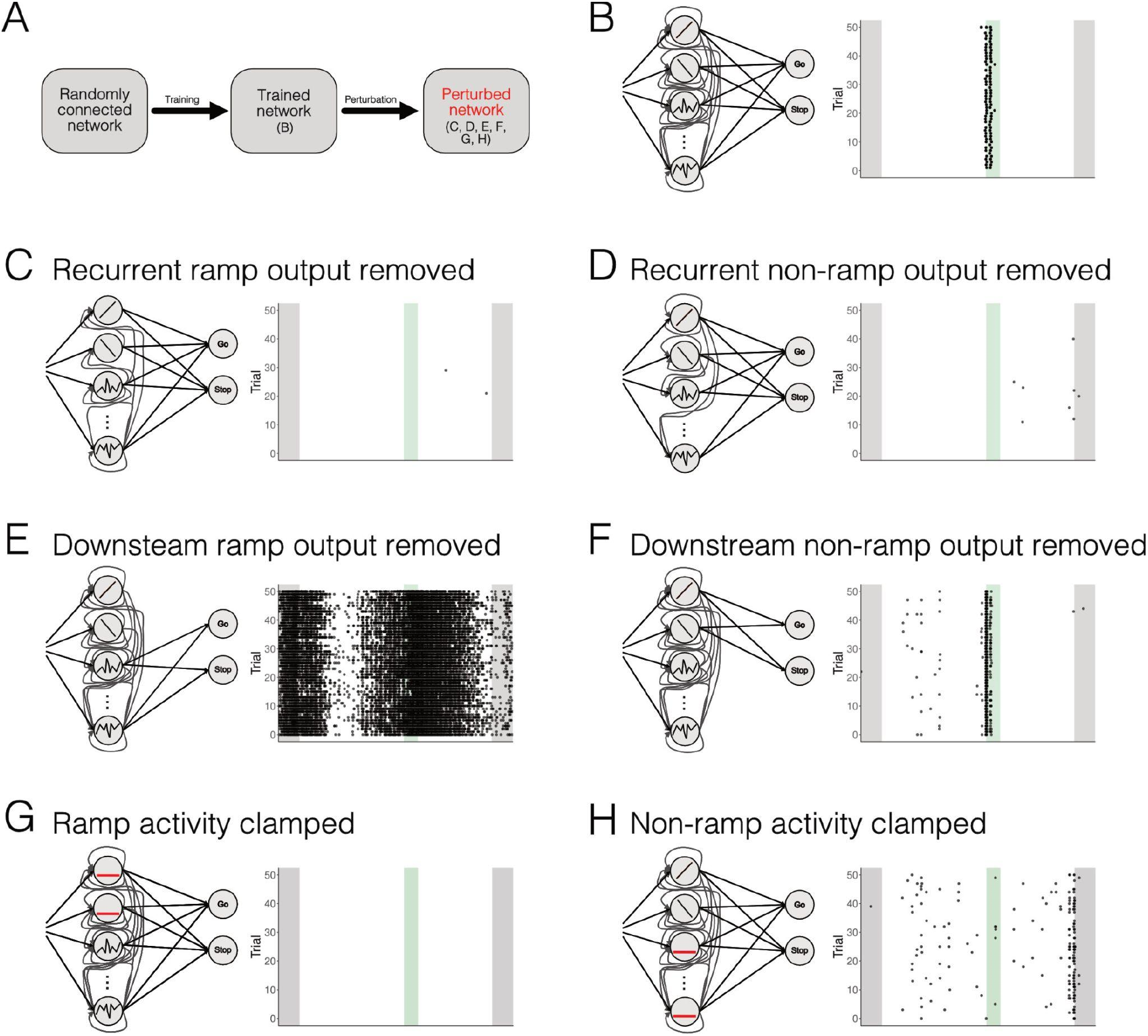
Perturbation of ramping activity disrupts control of spatial stopping by trained recurrent networks. (A) Perturbations were introduced after training randomly connected recurrent networks in the linear location estimation task. Raster plots show stopping locations of the trained network in control conditions (B), after blocking recurrent outputs from ramping neurons (C) and non-ramping neurons (D), after blocking the outputs to movement control neurons from ramping neurons (E) and non-ramping neurons (F), and after clamping the activity of ramping neurons (G) and non-ramping neurons (H) to its mean value. Criteria for selection of ramping and non-ramping neurons are in the methods.

Together, these analyses show that network configurations that solve the location estimation task are discoverable by reinforcement rather than being designed a priori, and that ramping activity is critical to the success of these network configurations. This is consistent with ramping activity patterns being able to play causal roles in recall of location memories.

## Discussion

Our results show that retrohippocampal neurons employ ramping neural activity to generate structured representations of learned environments, with reward location associated with switches in the ramp slope and resetting of the firing rate of ramping neurons. These representations could be maintained in the absence of location-specific cues, indicating that they result from recall of a spatial memory. As the direction of ramping activity could differ between segments of the behavioural track, representations of position may be specific to individual segments, potentially enabling ramp codes to combinatorially represent location and local context, for example by using ascending and descending ramp slopes to encode locations before and after the reward zone. These representations differ from the global metric provided by grid cells ^3,45^, but are reminiscent of robotic control strategies that segment the environment ^46^. The ramp-like firing within a track segment is also reminiscent of spatial firing of neurons in the ventral striatum in anticipation of rewarded locations ^26–32^, and of asymmetric firing of place cells in familiar track environments ^47^. However, retrohippocampal activity differs in the diverse combinations of firing rate slopes approaching and following the reward location (Figure 1), and in their insensitivity to whether reward is delivered (Figures 3 and 4). These distinct features of ramp-like firing of retrohippocampal neurons may be suited to stable representation of the structure of the learned environments and location within them relative to salient landmarks, which in the task here are the reward zone and the start and end of the track.

Our results extend the representational strategies through which neurons in the hippocampal formation might encode goals. In the hippocampus, learning about rewarded goal locations has been associated with an increase in the density of place fields ^19–22^, with the emergence of distinct discrete goal related firing fields ^20,23^, or when goal and reward locations are distinct with additional discrete firing fields at the goal location ^48–50^. In bats, discrete firing fields may also encode direction to goals and the distance from goals ^51^. In the medial entorhinal cortex, learning about rewarded goal location causes a shift in the location of nearby grid firing fields ^24,25^. In contrast, ramping activity that we describe here provides information about the distance to a goal, while resetting of ramps and changes in their polarity are associated with the goal location. Similar to discrete directional and distance codes in the bat hippocampus ^51^, ramping activity profiles did not require cues associated with the goal to be present. Thus, memory recall appears to be able to generate both types of representation.

What signals initiate, maintain and terminate ramping activity? Initiation of ramping activity was associated with the start of the virtual track suggesting that it must be determined by visual signals. The progression of ramping activity was then best explained by the animal’s track position alone or in conjunction with its speed and / or acceleration (Figure 2). Several observations indicate that for most neurons modulation of their firing rate by speed or acceleration is unlikely to be the primary drive of ramping activity. First, position rather than speed or acceleration was usually the dominant coefficient explaining their firing (Figure 2B). Second, pure speed and acceleration coding neurons were identified more frequently among non-ramping then ramping neurons (Figure 2C), and speed and acceleration profiles during the task differed from ramping activity profiles (Figure 2A). Third, evaluation of activity profiles on trials in which mice failed to stop for rewards enables further dissociation of the contributions of position, speed and acceleration (Figure 3). Despite the speed profile being reversed on these trials ramping profiles were in general maintained. Similar considerations argue that changes in speed or acceleration are unlikely to account for resetting of ramping activity or switches in ramp slope in the regions of the reward zone. An alternative possibility is that reward expectation signals rather than position per se drive ramping activity (cf. ^48–50^); in this scenario activity profiles on probe trials must still reflect recall of the structure of the track and task. While ramping activity on the track segment after the reward zone is inconsistent with reward expectation, dissociating reward expectation from location coding will require tasks in which the stopping location and reward delivery location differ (cf. ^48^).

According to current theories for spatial computation, the location task we use here could be solved using path integration by grid cells to drive sequential activation of place cells along the track. In this scenario visual cues anchor the grid representations to the start of the track and to the reward zone. However, it is challenging for neural circuits to read out metric representations of distance to the stopping position from discrete spatial firing fields without using graded intermediate representations ^10,11,17^. Our results are consistent with proposed solutions to this problem in which the structure of the environment is stored in the weights of synaptic inputs from place or grid cells onto ramping neurons ^10,11,17^. In this case, learning modifies synaptic weights so that the discrete spatial inputs from upstream neurons drive ramping activity profiles across the length of the track. Ramping activity within a track segment could also be generated without place or grid cell input through integration of velocity signals by recurrent connectivity (Figures 6-7) or intrinsic dynamics of entorhinal neurons ^52–55^, or could be driven by inputs from postrhinal neurons ^56^. The intrinsic dynamics of recurrent networks could also mediate interruption of ramping activity at reward locations (Figures 6-7). In this case, a key difference from models in which grid or place inputs drive ramp firing is that the recurrent network performs a path integration-like computation and the reward zone location is encoded by plasticity in the synaptic weights within the recurrent network.

Our analysis of artificial neural networks that solved the location memory task is consistent with the idea that ramping activity contributes to control of learned behaviours (Figures 6 and 7). The ramping activity emerged without a priori design to make the artificial networks resemble the retrohippocampal cortices, other than through the selection of an initially unstructured recurrent architecture. Thus, ramping codes may be a general solution for recalls of location in track-like environments in the sense that their emergence is not tied specifically to the initial details of circuit organisation. Further investigation will be required to establish whether this applies across other learning algorithms and to determine the extent to which the learned synaptic weights resemble connectivity in the retrohippocampal cortices. Because different spatial coding schemes emerge in artificial neural networks trained with different tasks (e.g. ^57,58^), it will also be of interest to explore how task contingencies interact with learning rules to shape emergent representations. Impaired spatial localisation after manipulation of the outputs from ramping neurons in the artificial networks suggests that ramping activity can play a necessary role in control of spatial behaviours. Similar tests of causality in animal models will require strategies for selective manipulation of ramping neurons during behaviours.

Given that deep layers of entorhinal cortex have extensive projections throughout the telencephalon ^59–61^, ramping activity may be used by motor control neurons receiving input from the retrohippocampal cortices, or as a template for ramping activity of neurons in striatal circuits that receive direct inputs from retrohippocampal neurons. Thus, our results suggest a model for memory recall in which a neural path integrator drives ramping neural activity that reconfigures at salient locations and that may be suitable for control of behaviours by downstream brain structures.

## Acknowledgements

We thank Richard Morris for feedback on an earlier version of the manuscript, Robert Wallace for assistance with micro-CT imaging, Holly Stevens for technical assistance, and Gülşen Sürmeli, Ian Duguid and members of the Sürmeli, Duguid and Nolan labs for helpful discussions. This work was supported by grants to M.F.N. from the Wellcome Trust (200855/Z/16/Z) and the BBSRC (BB/ L010496/1); by grants from the Simons Initiative for the Developing Brain to M.F.N; by a College of Medicine and Veterinary Medicine PhD Studentship, funded by the Thomas Work Fellowship, to K.G., by the Wellcome Trust (108890/Z/15/Z) Translational Neuroscience PhD programme to I.H. and by the Medical Research Council Precision Medicine PhD programme to H.C. This work made use of resources provided by the Edinburgh Compute and Data Facility. For the purpose of open access, the author has applied a CC BY public copyright licence to any Author Accepted Manuscript version arising from this submission.

## Author contributions

ST and MFN conceived of the study. ST, HC, IH, WKT, KZG, ERW and MFN contributed experimental design and analyses. ST, HC, IH, WKT, JH and WY acquired and analysed data. MN and ST drafted the manuscript. All authors contributed to editing of the manuscript.

## STAR Methods

### RESOURCE AVAILABILITY

#### Lead contact

Further information and requests for resources and reagents should be directed to and will be fulfilled by the lead contact: Matthew Nolan.

#### Materials availability

This study did not generate new unique reagents.

#### Data and code availability

Data will be deposited at https://datashare.ed.ac.uk/handle/10283/777 and will be publicly available as of the date of publication. DOIs will be listed in the key resources table. All original code used for analysis of the data is deposited at https://github.com/MattNolanLab and will be publicly available as of the date of publication. DOIs will be listed in the key resources table. Any additional information required to reanalyze the data reported in this paper is available from the lead contact upon request

### EXPERIMENTAL MODEL

Experiments used 11 male mcos wild type mice 7-14 weeks (mean = 12.6, SD = 0.71) old at the time of the implant surgery. Males were used for consistency in their size, which was important for operation of the behavioural apparatus. One week prior to surgery animals were group housed (3-5 mice per cage) in a holding room on a reversed 12 hour on/off light cycle (light from 7 pm to 7 am). After surgery mice were single housed in cages that contained a running wheel and cardboard igloo for additional enrichment. Prior to surgery and for 2-3 days following surgery mice were given standard laboratory chow and water *ad libitum*.

### METHOD DETAILS

All animal experiments were carried out under a project licence granted by the UK Home Office, were approved by the Animal Welfare and Ethical Review Board of the University of Edinburgh School of Medicine and Veterinary Medicine, and conformed with the UK Animals (Scientific Procedures) Act 1986 and the European Directive 86/609/EEC on the protection of animals used for experimental purposes.

#### Microdrive fabrication

Microdrive fabrication for in vivo electrophysiology recordings was similar to our previous work ^62^. 16-channel microdrives consisting of 4 tetrodes were built by threading 90% platinum, 10% iridium wire (18 μm HML-coated, Neuralynx) to an EIB-16 board (Neuralynx) via an inner cannula (21 gauge 9 mm long). The board was then covered in epoxy and once dried a poor lady frame (Axona) was cemented to the side. An outer cannula (17 gauge 7 mm), sanded at a 45 degree angle on one end to fit the curvature of the skull, was placed around the inner cannula. The outer cannula was secured temporarily using vaseline, allowing it to be lowered during the surgery. Tetrodes were then trimmed to ~3 mm using ceramic scissors (Science Tools, Germany) and gold plated (Non-cyanide gold plating solution, Neuralynx) using a custom built tetrode plater ^63^ until the impedance was between 150 and 200 kΩ.

#### Surgical procedures

Before surgery, tips of the tetrodes were washed with ethanol and then with sterile saline. General surgical procedures were carried out as described previously ^59^. Inhalation anaesthesia was induced using 5 % isoflurane / 95 % oxygen, and sustained at 1 – 2 % isoflurane / 98-99 % oxygen with a flow rate of 1 L / minute. An oval incision was made to expose the surface of the skull and a RIVETS head-post ^64^ attached to the skull and held in place with UV curing resin cement (RelyX™ Unicem, 3M). For electrical grounding, two small craniotomies were drilled on the left hemisphere ~3.4 mm lateral, and ~1 mm rostral relative to Bregma and the centre of the intraparietal plate respectively. M1 x 4 mm screws (AccuGroup SFE-M1-4-A2) were then implanted just below the surface of the skull and cemented in place.

The microdrive was attached to an Omnetics to Mill-Max adaptor (Axona, HSADPT-NN1), which was then attached to the stereotaxic frame. The tetrodes were lowered 1.2 - 1.4 mm into the brain, beginning at 3.4 mm lateral from Bregma (right hemisphere) and along the lambdoid suture, and at an angle of - 16° in order to target retrohippocampal cortices. The outer cannula was then lowered and sterile Vaseline carefully placed around the rim to seal the craniotomy. The implant was fixed to the skull with UV curing resin. After the resin hardened, the grounding wires were wrapped around the grounding screws and fixed with silver paint (RS components 101-5621). A layer of resin was applied to cover the grounding screws and any holes were filled with dental cement (Simplex Rapid). Mice recovered on a heat mat for ~20 minutes, and had unlimited access to Vetergesic jelly (0.5 mg / kg of body weight buprenorphine in raspberry jelly) for 12 hours after surgery. Mice were given a minimum of 2 days postoperative recovery before proceeding.

#### Handling and habituation

Handling and habituation of mice was similar to that described previously ^33,62^. Mice were handled twice a day for 5 - 10 minutes for one week following surgery. Animals were then habituated to the virtual reality setup for 10 and 20 minutes respectively over two consecutive days. After each habituation session animals were given access to soy milk in their cages to familiarise them with the reward and were given access to a larger arena for 5 - 10 minutes of free exploration.

#### Food restriction

To motivate mice to obtain rewards in the location memory task their access to food was restricted so that their body weight was maintained at ~85% of its baseline value ^33^. From 4-5 days before starting training an allotted food amount, determined on a daily basis to maintain the animal’s weight at its target value, was made available to the mice. The baseline weight of the animal was calculated from its weight prior to restriction and normalised to the expected daily growth for the animal’s age.

#### Experimental design overview

Experimental days involved two recording sessions; one in virtual reality and one in an open arena (Figure 5A). On these days animals were collected from the holding room 30 - 60 minutes before the first recording session. Before recording, animals were handled for 5 - 10 minutes, weighed and placed for 10-20 minutes in a cage filled with objects and a running wheel. Animals were then head fixed in the virtual reality setup and performed the location memory task for approximately 30 minutes. After the recording session, animals were placed back in the object-filled arena for 30 minutes before a second recording session in an open field arena for 30 minutes. They were then returned to their home cage. Tetrodes were typically lowered by 50 - 100 μm after each session. At the end of the experiment animals were sacrificed and the tissue processed for anatomical localisation of the recording location (described below). Animals in which the recording location was outside the retrohippocampal areas were excluded from analysis (n = 2).

#### Location memory task

Mice were trained to obtain rewards at a marked location on a virtual linear track as described previously ^33^. During recording sessions animals were head fixed in the virtual reality setup for approximately 30 minutes. Mice were head fixed using a Rivets clamp (Ronal Tool Company, Inc)^64^ and ran on a custom made cylindrical treadmill fitted with a rotary encoder (Pewatron). A feeding tube placed in front of the animal dispensed soy milk rewards (5 - 10 μl per reward). Virtual tracks were generated using Blender3D (blender.com). The track had length 2 m, with a 60 cm track zone, a 20 cm reward zone, a second 60 cm track zone and a 60 cm black box to separate successive trials. The walls and floor of the track were labelled with a repeating pattern (Figure 1A). On trials in which the reward zone was visible it was indicated by distinct vertical green and black bars. At any point on the track the visible distance ahead of the mouse was 40 cm.

Electrophysiological signals from the 16 channel microdrive were acquired using an Intan headstage connected via an SPI cable (Intan Technologies, RHD2000 6-ft (1.8 m) standard SPI interface cable) attached to an Open Ephys acquisition board ^65^. Signals were filtered using the board’s analogue high and low cut filter (2.5 Hz -7603.8 Hz). Behavioural variables including position, trial number and trial type were calculated in Blender3D at 60 Hz as described previously ^33^. Using Blender’s embedded Python interpreter (Python API), custom scripts wrote each variable (60 Hz, 12-bit precision) to a data acquisition (DAQ) microcontroller (Arduino Due) which in turn sent the signals to the OpenEphys acquisition board via HDMI.

#### Open arena exploration

The open arena, as described in ^62^, consisted of a metal box with a square floor area, removable metal walls, and a metal frame (Frame parts from Kanya UK, C01-1, C20-10, A33-12, B49-75, B48-75, A39-31, ALU3).

For motion and head-direction tracking, a camera (Logitech B525, 1280 x 720 pixels Webcam, RS components 795-0876) was attached to the frame of the open arena. To record motion alongside head-direction data, red and green polystyrene balls were attached to the sides of the headstage. A custom Bonsai script then tracked the polystyrene balls during recording sessions based on their colour ^66^. To synchronize the position and electrophysiology data, an LED was attached to the side of the open arena in the field of view of the camera. Pulses were sent to the LED via an Arduino (Arduino Uno) as well as to Open Ephys acquisition board via the I/O board. The pulses were separated by randomly generated intervals (20 to 60 seconds) so that the two datastreams could be reconciled during subsequent analysis ^66^.

Electrophysiological signals were recorded using the custom made 16 channel microdrive described above. An Intan recording headstage with an SPI cable (Intan Technologies, RHD2000 6-ft (1.8 m) Ultra Thin SPI interface cable C3216) attached was used to connect the microdrive to a commutator (SPI cable adapter board, Intan Technologies C3430 and 3D printed holder, custom designed by Patrick Spooner). The commutator was connected to an Open Ephys acquisition board ^65^ that sent the electrophysiology signal to a computer where the signal was filtered between 2.5 Hz - 7603.8 Hz and displayed using the OpenEphys GUI.

#### Post recording anatomical assessment of tetrode location

To determine tetrode locations we used either Cresyl violet staining (n = 3 mice) or microcomputed tomography (microCT) (n = 7 mice) (Supplemental Data 1). Two mice that underwent cresyl violet staining had no visible tetrode tracks and are omitted from Supplemental Data 1. For these mice tetrode location was inferred by the presence of theta modulation of the local field potential and the spiking activity of recorded neurons (Supplemental Figure 3). For both methods mice were anaesthetised with isoflurane and given a lethal dose of sodium pentobarbital (Euthatal, Meridal Animal Health, UK).

For cresyl violet staining, we made an electrolytic lesion using 2 x 2 ms, 20 μA current pulses per tetrode. The mice were fixed by transcardial perfusion of ice cold phosphate buffered saline (PBS; Invitrogen) followed by 4 % paraformaldehyde (PFA; Sigma Aldrich) in 0.1 M phosphate buffer (PB; Sigma Aldrich). Brains were removed, left in 4 % PFA in 0.1 M PB overnight and then in 30 % sucrose in PBS for two nights. A freezing microtome was used to cut 50 μm sagittal slices, which were mounted on polarised slides, dried overnight and then placed in xylene before rehydration in varying concentrations of ethanol (100%, 90% then 70% for 1-2 minutes each). Once washed, sections were placed in Cresyl violet for 2 mins before additional washing and dehydration (70% then 90% for 1.5 minutes each followed by two 30 second washes in 100%). Sections were then placed in xylene and mounted with Dibutylphthalate Polystyrene Xylene (DPX). Imaging of sections was performed with a slide scanner (Zeiss Axioscan.Z1). Sections were then manually inspected for the lesion site.

For microCT imaging, we modified a protocol used previously for imaging rat brains ^67^. Animals were perfused with a mixture of PFA and glutaraldehyde and then left in the same solution for 2 nights. Brains were then washed in ddH20 before being incubated at 4° for two weeks in 2 % osmium tetroxide (2% OsO4). After osmium staining brains were washed in ddH20, dehydrated in ethanol and then embedded in resin. Once the resin had cured, we placed the sample in a micro-CT scanner (Skyscan 1172, Bruker, Kontich, Belgium) which generated a 3D image of the sample. Scanning parameters were: source voltage 54 kV, current 185 μA, exposure 885 ms with a 0.5 mm aluminium filter between the X-ray source and the sample ^68^. After acquiring the scan the dataset was reconstructed (CTan software, v1.13.5.1) and viewed with DataViewer (Bruker).

For the Cresyl violet and microCT methods, tetrodes were localised relative to landmarks in version 2 of the Allen Reference Atlas for the mouse brain (https://mouse.brain-map.org/static/atlas). Tetrodes were assigned either to the MEC (n = 5 mice), the pre/parasubiculum (n = 2), or when the location was ambiguous between these areas to retrohippocampal cortex (n = 3)(Supplemental Data 1). Theta frequency field potentials and theta phase locking of spike firing (Supplemental Figure 3) were consistent with previous reports for these regions ^40,69–71^. We did not attempt to differentiate between presubiculum and parasubiculum as in the mouse their borders are not well defined without additional markers and there is disagreement between reference atlases when using landmarks as reference points. Location of all tetrodes in the retrohippocampal region, defined as including the MEC, presubiculum and parasubiculum (e.g. see ^72^) was corroborated by electrophysiological recording of a theta field potential and theta modulation of the firing of neurons (cf. ^40^)(see Supplemental Figure 5 and analyses section below). In contrast, neurons in nearby structures such as the postrhinal cortex do not show theta phase locked firing ^56^. The animals from which we were unable to localise the tetrodes anatomically, but in which field potential recordings and spike firing showed theta modulation, were assigned to the retrohippocampal group.

### QUANTIFICATION AND STATISTICAL ANALYSES

Analyses were carried out using Python (version 3.5.1) and R versions: 3.3.1 to 4.2.0.

#### Spike sorting

Spikes were isolated from electrophysiological data using an automated pipeline based on MountainSort (v 0.11.5 and dependencies)^73^ and similar to that described in ^62^. Data pre-processing steps converted Open Ephys files to mda format, filtered signals between 600 Hz - 6000 Hz and then performed spatial whitening over all channels. Events were detected from peaks > 3 standard deviations above baseline and separated by at least 0.33 ms from other events on the same channel. A 10-dimensional feature space was generated using the first 10 principal components of the detected waveforms and spike sorting using the ISO SPLIT algorithm performed on the feature space ^73^. Cluster quality was evaluated using metrics for isolation, noise-overlap, and peak signal to noise ratio ^73^. Units that had a firing rate > 0.5 Hz, isolation > 0.9, noise overlap < 0.05, and peak signal to noise ratio > 1 were accepted for further analysis.

#### Classification of ramp-like activity

To classify ramping neurons (Figure 1, Supplemental Figure 4) we generated firing rate maps for each trial by binning the track into 1 cm bins, summing the number of spikes in each bin and dividing by the time the animal spent there. Bins in which the animal’s speed was < 3 cm/s were removed. For the purposes of data visualisation, firing rates were smoothed by convolution with a Gaussian filter (SD = 2 cm, SciPy Python package ^74^) and then averaged across trials. For each neuron, we also generated 1000 shuffled datasets by adding a random time drawn from a uniform distribution between 20-580 seconds and added this to the timestamp of each spike. Random times were drawn independently for each trial and shuffle. Spike times that exceeded the recording length were wrapped around to the start of the session. Spike locations were recomputed from the shuffled spike times and a firing rate map generated as described previously. We then fit linear models to firing rate as a function of position across the regions of the track before (0-60 cm) or after the reward zone (80-140 cm). This was done for each of the 1000 shuffled firing rates and the measured dataset for each neuron. The significance (p) values obtained from fitting the linear models were corrected for multiple comparisons using the Benjamini & Hochberg method ^75^. We classified neurons as having ramp-like activity with positive (or negative) slopes if the linear model was significant (corrected p < 0.01) and the coefficients of the model were outside the 5 - 95 % intervals of the shuffled data. Neurons not meeting these criteria were allocated to the unclassified group. We focussed our analyses on six groups of neurons separated according to their slope classifications: positive before the reward zone and after the reward zone (+ +), positive before the reward zone and negative after (+ -), positive before the reward zone and unclassified after (+ un), negative before the reward zone and negative after (- -), negative before the reward zone and positive after (- +) or negative before the reward zone and unclassified after (- un).

#### Ramp score calculation

As a complementary method to identify ramp-like firing we calculated a ramp score based on the correlation between firing rate and the animal’s position (Supplemental Figure 2). This ramp score is related to the score used previously for identification of speed cells ^76^. For each cell we calculated separate ramp scores for the track segments before and after the reward zone (Supplemental Figure 2A).

We first smoothed the mean firing rate of each cell in each binned position using a Savitzky-Golay filter (window size: 7, polynomial order: 1) and then identified locations before the reward zone at which peaks and troughs in the firing rate were present by the change of the sign of the slope. Using all turning points (peaks and troughs) in the smoothed curve, together with two endpoints (i.e. beginning and end) of the track segment before the reward zone, we construct a set of potential track segments. For example, if there is a single peak at 30 cm, with two endpoints of the zone at 0 cm and 60 cm, then the set of segments will be (0, 30), (30, 60), (0, 60), where each segment is defined by its starting and ending locations.

To avoid bias from location bins with many data points (e.g. when the animal slowed down in the reward zone), we oversample the original unsmoothed firing rate so that all segments have the same number of points. Then in each of the track segments *d_i_*, using the balanced firing rate *fr*, we calculate the Pearson correlation coefficient *r_i_* between the firing rate and the location *loc*.

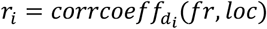

To calculate the ramp score for each segment we first determine the ratio of the length of the track segment to the whole track,

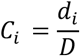

where *d_i_* is the length of the i-th track segment, *D* is the length of the track region of interest (e.g. the regions before or after the reward zone). The ramp score *S_i_* of this segment is then the harmonic mean of the correlation coefficient and the segment ratio:

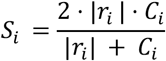

A harmonic mean is used to ensure that a high ramp score will mean that the cell has a high correlation coefficient and a high segment ratio.

To determine the ramp score for each cell, the ramp scores of all possible segments were calculated and the final ramp score *S_R_* of the cell is chosen as the ramp score that has the largest magnitude. The sign of the ramp score indicates whether the slope is positive or negative.

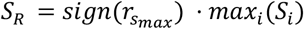

where *r_s_max__* is the correlation coefficient of the track segment corresponding to the maximum ramp score.

#### Generalised linear mixed effect models for position, speed and acceleration

To quantify the influence of position, speed and acceleration on the firing rate of each neuron, we built generalised linear mixed effect models to predict firing rate using position, speed and acceleration as fixed effects and trial number as a random effect. For each neuron, all variables were binned into 100 ms blocks and bins when the animal’s speed was < 3 cm/s were removed ^77,78^. For the purposes of data visualisation, but not model fitting, firing rates and kinematic variables were smoothed by convolution with a Gaussian filter (SD = 200 ms, SciPy Python package ^74^). The average speed and acceleration of the animal across the whole recording was calculated, and bins in which speed or acceleration was 3 standard deviations (SDs) above the average for that variable were removed. A Poisson model with the configuration ‘Rate ~ Position + Speed + Acceleration + (1 + Position | Trials)’ and log linker function was fit using the lme4 package (version 1.1-12) in R ^79^. The ANOVA function provided by the car package (version 3.0-9) was used to calculate significance values for each model coefficient ^80^. To classify neurons as being modulated by position (P), speed (S), acceleration (A), or a combination thereof, we used a significance threshold of 0.01. For example, a neuron with p < 0.01 for position and acceleration and ≥ 0.01 for speed coefficients was classified as a PA neuron. P values obtained from the ANOVA were corrected for multiple comparisons using the Benjamini & Hochberg method ^75^.

#### Classification of trial outcomes

Trials were classified by task performance into hits and misses based on whether the mouse registered a stop (speed < 4.7 cm/s) within the reward zone ^33^ or not. Miss trials were split into near-hits (tries) and run-throughs (runs), by comparing their average speeds in the reward zone to hit trials. The 95th percentile of average speeds in the reward zone for hit trials in a single session was used to discriminate between tries (< 95th percentile speed) and run trials (> 95th percentile speed).

#### Calculation of average speed

The average speed as a function of location on the track for each session was calculated by binning speed into 1 cm location bins for each trial and then smoothing by convolution with a Gaussian filter (SD = 2 cm, SciPy Python package ^74^). Speed for each location bin was then averaged over trials. The mean speed across all sessions was calculated for each animal for each trial outcome. The mean and standard error across all animals was calculated from the means for each animal.

#### Calculation of spiking theta index

Theta index was calculated following methods in ^71^ (Supplemental Figure 3A). Spike times were transformed into instantaneous firing rates. A power spectrum was then estimated using Welch’s method. The theta index was defined as

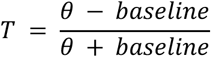

where θ is the mean theta band power (6-10 Hz) and baseline is the mean power of two adjacent frequency bands (3-5 and 11-13 Hz).

#### Power spectral analyses of local field potentials

We computed the power spectra across recording sessions for each channel for all raw voltage samples where the mouse was moving (> 3 cm/s). Power was estimated using Welch’s method using a 2 second (60000 samples at 30Hz sample rate) Hanning window and averaged across channels.

#### Spike half width calculation

Spike half width (the width of the spike at half-maximal amplitude) for each neuron was calculated as described in ^81^. For each neuron 50 spike waveforms were randomly selected and an average waveform generated. Spike amplitude was defined as the difference between the baseline voltage, calculated as the lowest value, and the positive voltage peak, calculated as the highest value. Spike half width was then calculated as the time (ms) the voltage stayed above half the maximal spike amplitude.

#### Classification of putative inhibitory and excitatory cells

Putative interneurons were defined as neurons with mean firing rate > 10 Hz, similar to previous studies in entorhinal populations ^82^ (Supplemental 7C). Excitatory neurons were classified as mean firing rates < 10 Hz. Consistent with this, spike half width in cells classified as putative interneurons were typically shorter than cells classified as excitatory (Supplemental 7C).

#### Classification of cell types in the open arena

Neurons were identified across recording sessions in the location memory task and open arena by concatenating the raw voltage channels from each session and then spike sorting. To classify neurons based on their activity in the open arena we followed methods in Diehl et al. (2017) to assign cells into one of five hierarchically organised functional cell types: grid > border > other spatial > head direction > non-spatial. Neurons were assigned to the highest cell type in the hierarchy for which their corresponding identity metric was greater than the 99th percentile of the same scores from 1000 shuffled datasets. For example, a cell with grid score above the 99th percentile was classified as a grid cell irrespective of scores in other categories. To generate shuffled spike data, we drew a single value from a uniform distribution between 20-580 seconds and added this to the timestamp of each spike. Spike times that exceeded the recording length were wrapped around to the start of the session. Spike locations were recomputed from the shuffled spike times and spatial scores calculated similar to measured data. Using this method we generated 1000 shuffles per cell and used the 99^th^ percentile of the scores as a threshold for belonging to the five cell types.

We used established scores for grid, border, head direction and spatial stability ^37,39,83^. Grid scores were defined as the difference between the minimum correlation coefficient for rate map autocorrelogram rotations of 60 and 120 degrees and the maximum correlation coefficient for autocorrelogram rotations of 30, 90 and 150 degrees (^37,39,83^). The firing rate map was calculated by binning spikes into 2.5 cm bins and dividing by the total time occupied in each bin and then smoothed with a Gaussian kernel. Autocorrelograms were calculated by sliding the rate map over all x and y bins and calculating a correlation score. Fields were detected in this autocorrelogram by converting it into a binary array using 20% of the maximal correlations as a threshold. If the binary array had more than 7 local maxima, a grid score was calculated. Correlations between the rotated autocorrelograms were then calculated using only a ring containing the 6 local maxima closest to the centre of the binary array and excluding the maximum at the centre. The ring was detected based on the average distance of the 6 fields near the centre of the autocorrelogram (middle border = 1.25* average distance, outer border = 0.25 * average distance).

To calculate border scores (^37,39,83^), firing fields were identified from the firing rate map by collecting neighbouring bins with firing rates greater than 30% the maximal firing rate and covering at least 200 cm^2^. For each putative field, the mean firing distance dm was computed as the average distance to the wall for each bin, divided by the mean firing rate of the field. To obtain a value between 0 and 1, dm was normalised by the shortest distance from the centre of the arena to the wall. The maximal coverage cm of a field was measured as the maximum proportion of field bins occupying any of the four edges of the firing rate map. The border score was then given by

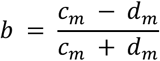

To calculate head direction scores (^37,39,83^), the head direction recorded at times corresponding to firing events were binned into 360 bins between 0 and 2π and was normalised by the duration of time spent occupying each directional bin. This polar histogram was then smoothed with a rolling sum with a window size of 23 degrees. The head direction score is the length of the mean vector of this polar histogram. To obtain this length, the x and y components are computed by first calculating dx and dy in a unit circle (radius = 1) in steps of 1 degree as

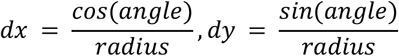

The head direction score is then given using the Pythagorean theorem by

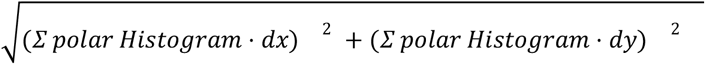

For assessing spatial stability, we used two separate metrics, spatial information and the within-session spatial correlation to identify spatial cells (^37–41^). We calculated the spatial information per spike as

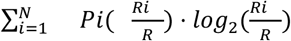

where i indexes a position bin in the firing rate map, N is the number of bins, Pi is the occupancy probability, Ri is the firing rate in the bin, R is the mean firing rate. The within-session spatial correlation is calculated by computing the Pearson correlation between the firing rate map computed from the first half session and the second half session. Bins that were not visited in both halves were excluded from the calculation.

#### Neural network model

We use a reinforcement learning algorithm to train artificial neural networks to solve a task similar to that used for the neuronal recordings. In the artificial task, the track consists of 1010 to 1150 locations (depending on track randomisation), each with a corresponding image. The track layout was similar to the experimental track, but differed in that visual distance to the reward zone was increased to make the task more challenging and the visual cues indicating the reward zone location were removed in order to promote learning of cue-independent behavioural strategies. The reinforcement learning agent could either choose to take a step forward or to stop. If the agent chooses to move forwards, the image corresponding to the next location in the track is shown to the network. The optimal behaviour is to keep moving forwards to the hidden reward zone, stop once to acquire the reward, and start moving to the end of the track to proceed to the next trial.

We cropped and re-sampled images used for the track in the recording experiment to 89×89 pixels and used them as the input to the artificial network. Each image frame of the track was first processed by a three-layer convolutional neural network before being fed into a discrete-time recurrent neural network (RNN) ^84^. A two-unit linear layer was used to read out the stop/go decision from the RNN. The recurrent states were reset after every 10 trials (which make up a training episode). To make the RNN outputs positive and comparable to neural firing rates, we chose the rectified linear unit (ReLU) as the activation function.

To simulate trial-to-trial variability in movement speed observed in the experimental data, the speed at which the agent moves was randomised at the start of each trial. For example, on ‘slow’ trials, the agent moved forwards 3 locations every time they chose to step forwards, but on ‘fast’ trials they moved 7 locations. The possible trial speeds were 3, 4, 5, 6, and 7. To simulate impreciseness of motor control, the number of locations moved forwards was also noisy, for example on the fastest trials there is an 80% chance of moving 7 locations forwards, with 15% probability for moving 7 or 9 (± 1) locations forwards, and 5% probability for 10 or 6 (± 2) locations forwards. Each black box zone was randomly padded by 0 to 70 locations. The artificial track was made up of repeated sequences of images, meaning it is not possible to know the true location on the track given a single image. These factors also decouple time from location so that the agent adopts a path integration strategy.

The task given to the agent differed from the task used for experimental recordings in that there was no visual reward zone cue. Training trials were equivalent to ‘non-beaconed trials’ and included a visual reward indicator in the form of a grey bar that was overlaid at the top of the image input for 5 timesteps if the agent stopped in the reward zone for the first time on a trial. This was designed to be analogous to the immediate feedback the experimental animals receive from a tone and soy milk reward. Probe trials, in which the indicator and reward were absent, were used only for evaluation of the network.

To train the agent to perform the task we used the proximal policy optimisation algorithm ^44^, which learned an action and value function. Documentation of the algorithm, including pseudocode, is available here: https://spinningup.openai.com/en/latest/algorithms/ppo.html. During training, stopping in the reward zone for the first time on a given trial resulted in a reward of +100, whereas any other stops resulted in a reward of -1. Moving forwards was a neutral action and had a reward of 0. These rewards were totalled for each trial. The network weights were updated using stochastic-gradient descent to maximise the reward ^85^. To find the optimal hyperparameters for training (such as learning rate), we performed random parameter searches using the Edinburgh Compute and Data Facility (ECDF) ^86^, using PyTorch 1.2.0 and cuDNN 10.1.105 ^87^ and adapted an existing Python implementation of the proximal policy optimisation algorithm ^88^. The hyperparameters used are in Supplemental Table 1.

## Supplemental figures

**Supplemental Figure 1.**
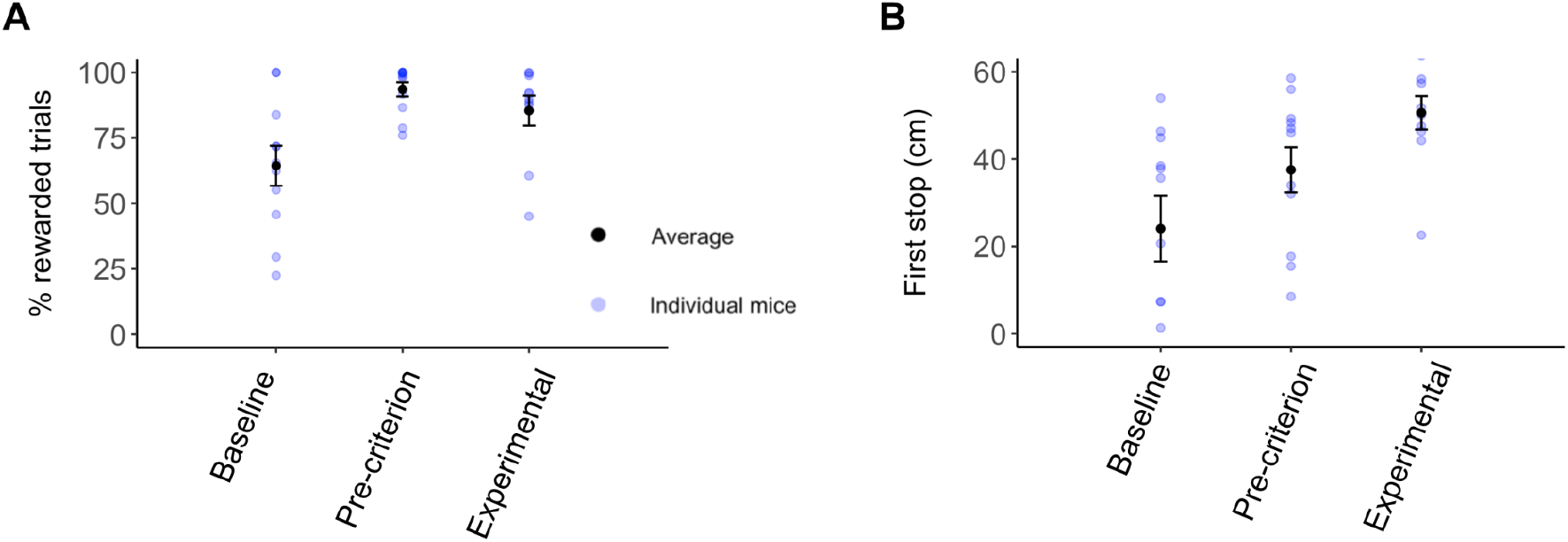
Behavioural training. (A) The percentage of rewarded trials for each mouse (blue) and averaged across mice (black, ± SEM) for the first training day (Baseline), the two consecutive days on which mice reached the criterion for introduction of probe trials (Pre-criterion) and the days after the criterion was passed (Experimental). Mice took 13.3 ± 7.1 days to reach the performance criterion. The reward rate depended on training period (Baseline, Pre-criterion or Experimental, p < 6.72 x 10^-7^, f =15.27; ANOVA). Compared with baseline sessions, the percentage of rewarded trials was larger during pre-criterion (adj p < 4 x 10^-7^, Tukey’s test) and experimental sessions (adj p < 3 x 10^-5^), whereas pre-criterion and experimental sessions did not differ significantly (adj p < 6 x 10^-2^). (B) The mean location at which mice first stopped on each trial for each mouse (blue) and averaged across all mice (black, ± SEM). Categories are as in (A). The first stop location depended on training period (Baseline, Pre-criterion or Experimental, p < 1.63 x 10^-15^, f = 40.6; ANOVA). Consistent with animals adopting a spatial stopping strategy ^33^, first stop locations were further along the track during experimental sessions compared with baseline (adj p < 1 x 10^-10^ Tukey’s test) and pre-criterion sessions (adj p < 1 x 10^-10^ Tukey’s test).

**Supplemental Figure 2.**
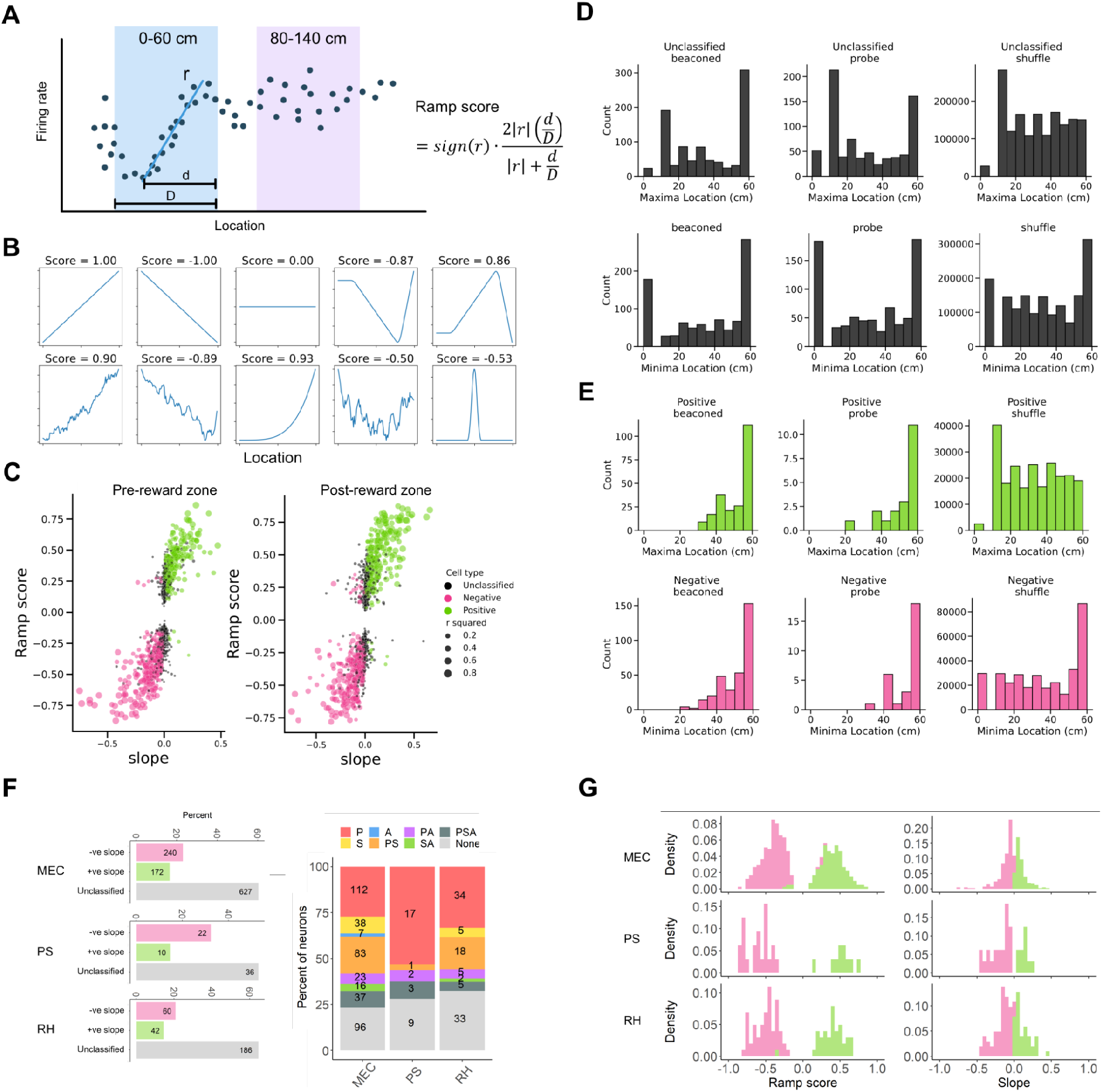
Evaluation of ramping activity using ‘ramp scores’. (A) Schematics showing the calculation of the ramp score. *r* is the correlation coefficient between the firing rate and the location. *d* is the length of the span of the ramp-like region. *D* is the length of the track region under consideration. Coloured regions indicate the segments of the track used for analysis. (B) Examples of ramp scores calculated for idealised reference waveforms. (C) Ramp scores plotted as a function of the slope of linear fits of firing rate to location for all neurons. (D-E) Locations of maxima and minima for unclassified neurons in the track segment before the reward zone (D) and neurons classified as having positive or negative slopes in the track segment before the reward zone (E). (F) Proportions of cells classified as having positive or negative slopes in the regions of the track before (left) and after (right) the reward zone (see Supplemental Figure 4) according to their locations in the medial entorhinal cortex (MEC), pre/parasubiculum (PS) or retrohippocampal cortex (RH). RH is used for neurons for which anatomical location in either MEC or PS could not be accurately determined, among these neurons the proportions classified as having firing that correlates with position (P), speed (S) and acceleration (A) (see Figure 2). (G) Distribution of slopes and ramp scores for MEC, PS and RH neurons that were classified as having positive or negative slopes.

**Supplemental Figure 3.**
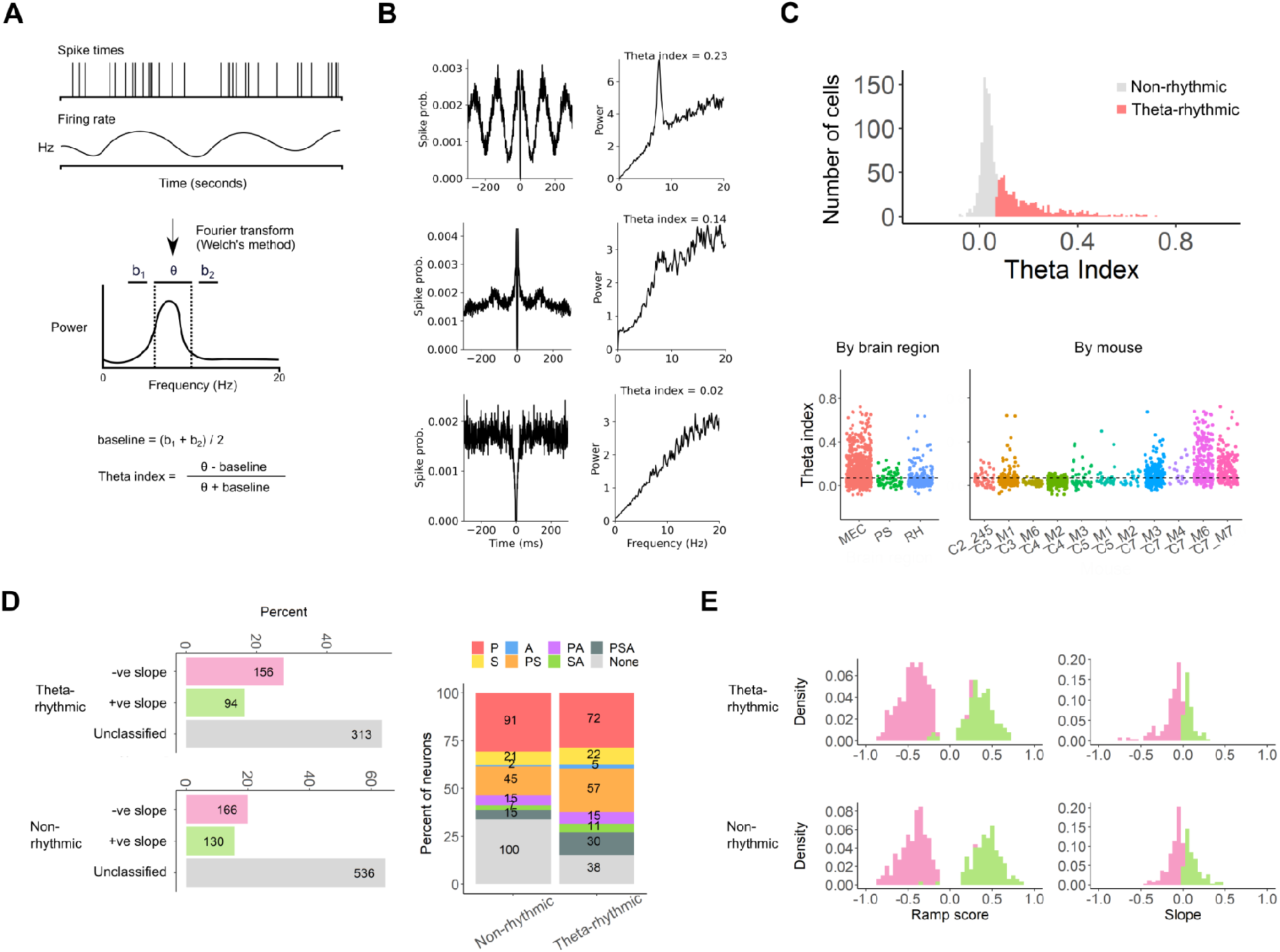
Theta modulation of neuronal firing. (A) Schematic illustrating the procedure for calculation of the theta index (see ^71^). Firing indices were transformed into instantaneous firing rates from which a power spectrum was estimated using Welch’s method. The theta index was defined as (θ-baseline)/(θ+baseline), where θ is the mean power in the theta band (6-10 Hz) and baseline (b) is the mean power of two adjacent frequency bands 3-5 Hz (b1) and 11-13 Hz (b2). (B) Autocorrelograms (left) and power spectra (right) for three simultaneously recorded neurons that exemplify strong (top), moderate (middle) and low (bottom) theta indices. (C) Histogram of the theta index across all cells (top) and scatter plots showing the theta index grouped by tetrode location (bottom left) and mouse (bottom right). Cells were classified as having theta-rhythmic firing (red in the histogram) when their theta index was > 0.07 (see ^71^). (D) Proportions of non-rhythmic (NR) cells and theta rhythmic (TR) cells classified as having positive or negative slopes (see Supplemental Figure 4) and among these neurons the proportions classified as having firing that correlates with position (P), speed (S) and acceleration (A) (see Figure 2). (E) Distribution of slopes and ramp scores for TR and NR neurons that were classified as having positive or negative slopes.

**Supplemental Figure 4.**
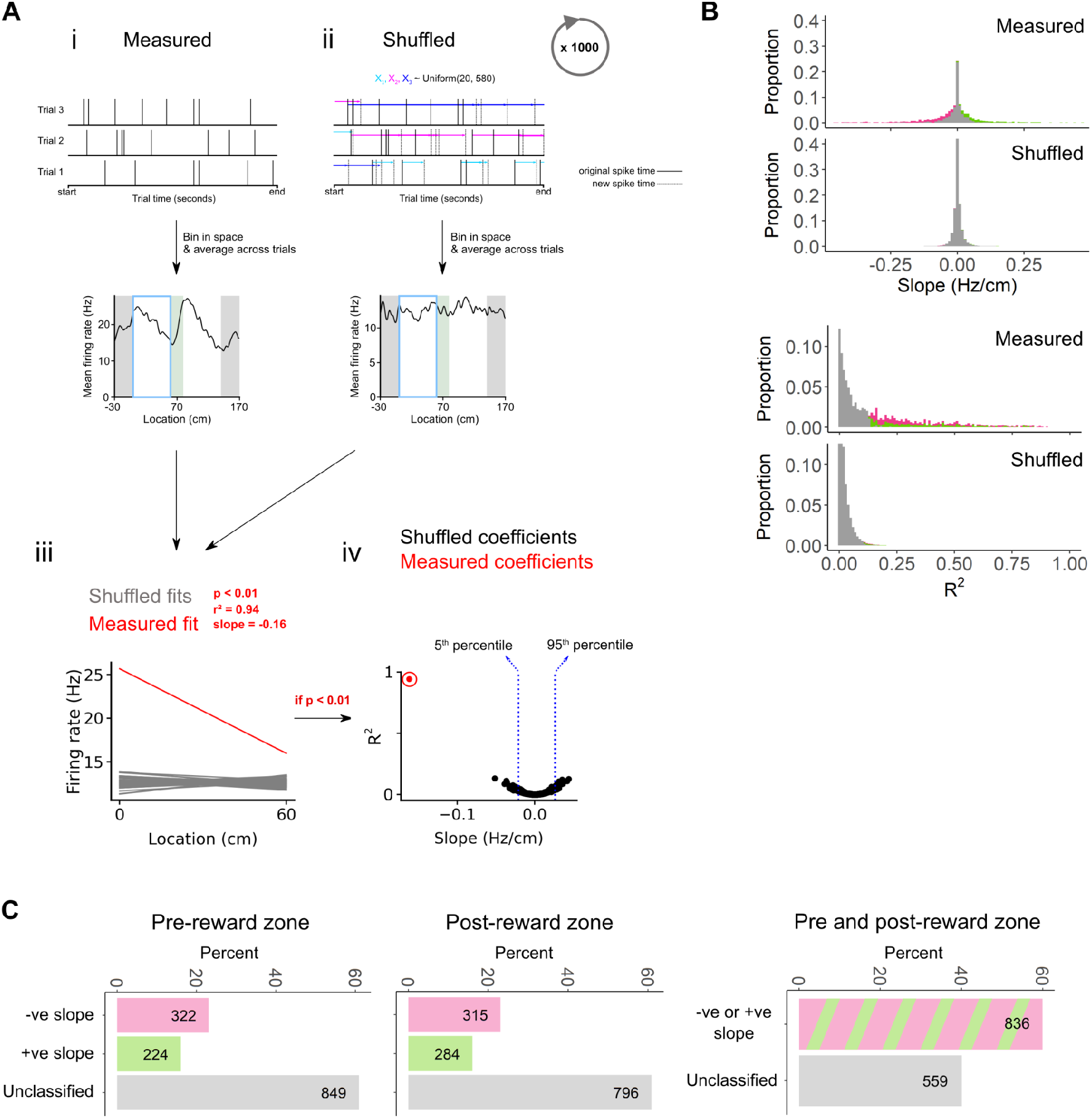
Quantification of ramp-like activity in shuffled data. (A) Shuffling procedure. Recorded (i) and a single set of shuffled firing rates (ii) plotted as a function of location (iii). Linear fits were calculated for the average recorded firing rate as a function of position on the track segment before the reward zone (0-60 cm, blue bar) and for the corresponding shuffled data (1000 shuffled datasets per neuron) (iv). Neurons were classified as ramping if the fit was significant (p < 0.01), and as having a positive ramp if the slope of the fit to their recorded data was above the 95^th^ percentile (blue dotted line) of slopes in the shuffled dataset (v) and a negative ramp if the slope was below the 5^th^ percentile. (B) The distribution of slopes (upper) and R^2^ (lower) of the model fit for each neuron. Neurons are colour-coded as having a positive slope (green), negative slope (pink) or unclassified (grey). Distribution of slope and R^2^ of the model fit for each shuffled dataset (n=1497501) is plotted in black below the corresponding experimental data. (C) Classification of experimentally recorded neurons based on their activity on the track segment before the reward zone(left), after the reward zone (middle), and both before and after the reward zone (right).

**Supplemental Figure 5.**
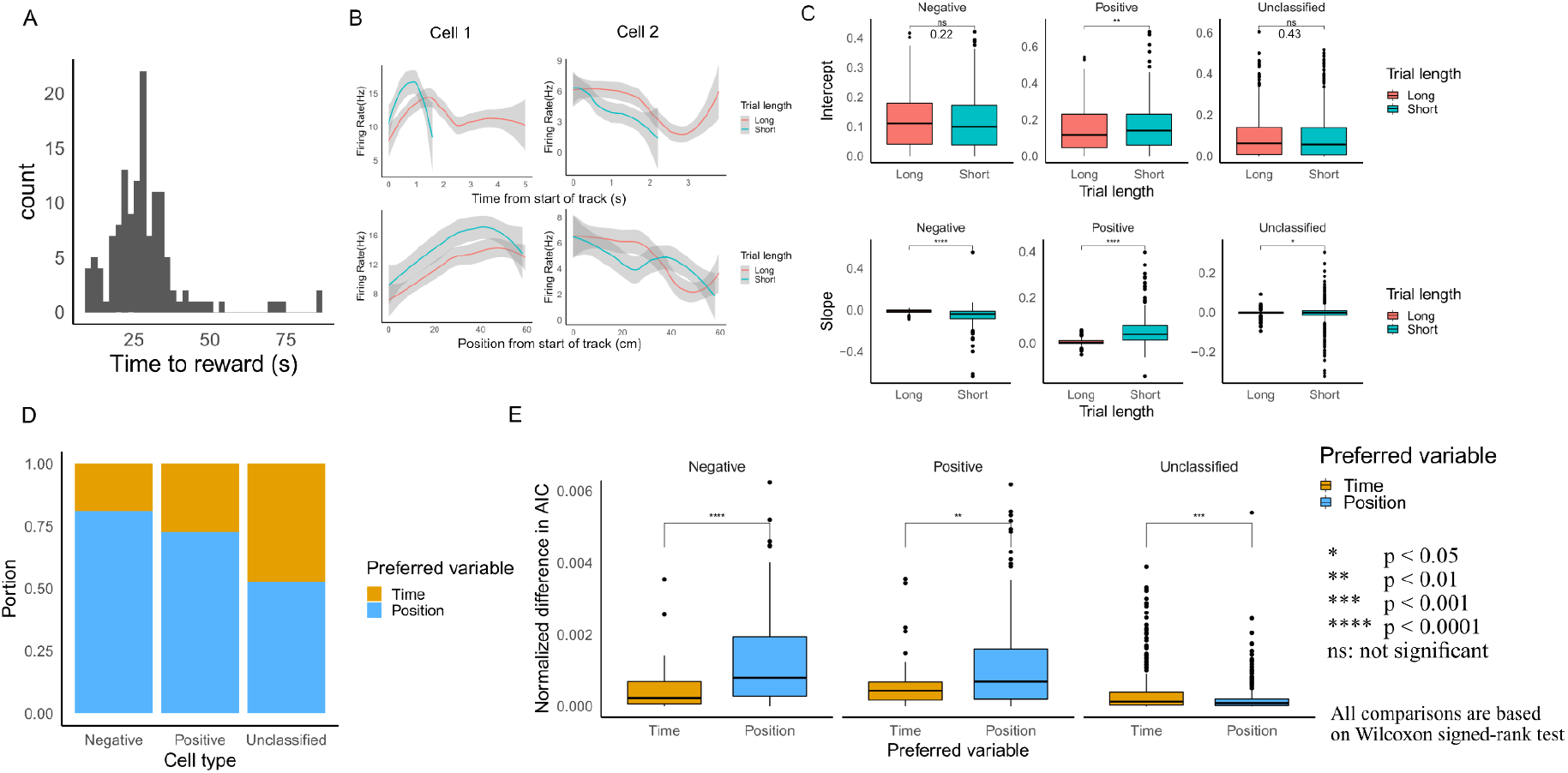
Evaluation of time versus position encoding. (A) Distribution of time to reach the reward zone in a representative session. (B) Mean firing rate of two ramping neurons in the track segment approaching the reward zone (track locations 0-60 cm) for short and long trials plotted as a function of time (upper) or position on the track (lower), from the same behavioural session as (A). Short trials are trials with duration less than the 25th percentile of all the trials in a session, while long trials are those with duration greater than the 75th percentile. A similar relationship between firing rate and distance on long and short trials is consistent with position encoding, whereas a similar relationship between firing rate and time is consistent with time encoding. (C) Comparison of the intercept (upper) and slope (lower) on short and long trials, for cells classified as having positive or negative slopes, when firing rate on the track segment approaching the reward zone is fit as a function of time. Significant differences between the slopes of short versus long trials are inconsistent with neurons encoding time (paired Wilcoxon signed-rank test). Values for significance estimates are shown in the figure. Test statistic top row, left to right: V=25072, V=9676, V=118384, bottom row: V=46875, V=2196, V=184201) (D) Proportion of cells encoding time or position based on comparison of Akaike Information Criterion (AIC) values obtained from mixed effect models fit to firing rate with either time or position as the fixed effect and trial number as the random effect. The model with the more negative AIC value was assumed to provide a better account of the data. In general the activity of neurons classified as having positive or negative slopes in the track segment following the reward zone were better predicted by models in which position rather than time is a fixed effect. (E) Comparison of the normalized difference in AIC between the position model and time model for different cell types. The difference is normalized by the sum of the AIC of position and time model. A larger difference is indicative of a model providing a better explanation for the data. Whereas position coding models were often strongly favoured, time encoding models were not (independent Wilcoxon signed-rank test, p-value for significance estimates are shown in the figure. Test statistic left to right: W = 4481, W = 3862, W = 108116).

**Supplemental Figure 6.**
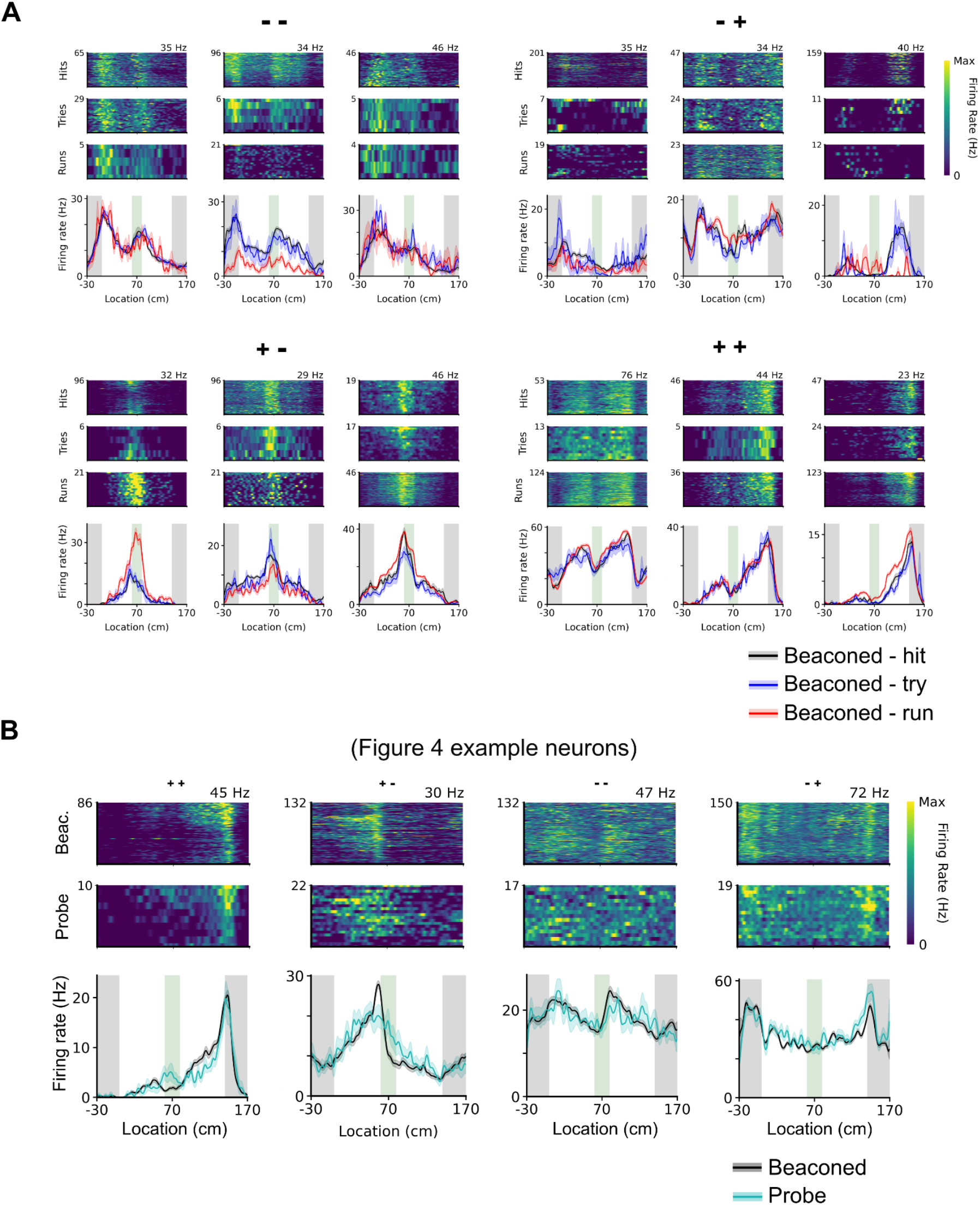
Examples of firing rate firing rate profiles for different trial outcomes and trial types. (A) Heat maps of firing rates (upper) and mean firing rate (lower) as a function of track location along the for all hit (left), try (centre) and run (right) trials for pairs of example neurons with track-wide ramp profiles categorised as ++, +-, -- and -+. The upper neuron of each pair is representative of neurons with firing rate profiles that are independent of trial outcome, while the lower neurons are representative of neurons in which the mean firing rate profiles appear to be modulated by the trial outcome. (B) Heat maps of firing rates (upper) and mean firing rate (lower) as a function of track location along the for probe and beacon trials for example neurons with track-wide ramp profiles categorised as ++, +-, -- and -+.

**Supplemental Figure 7.**
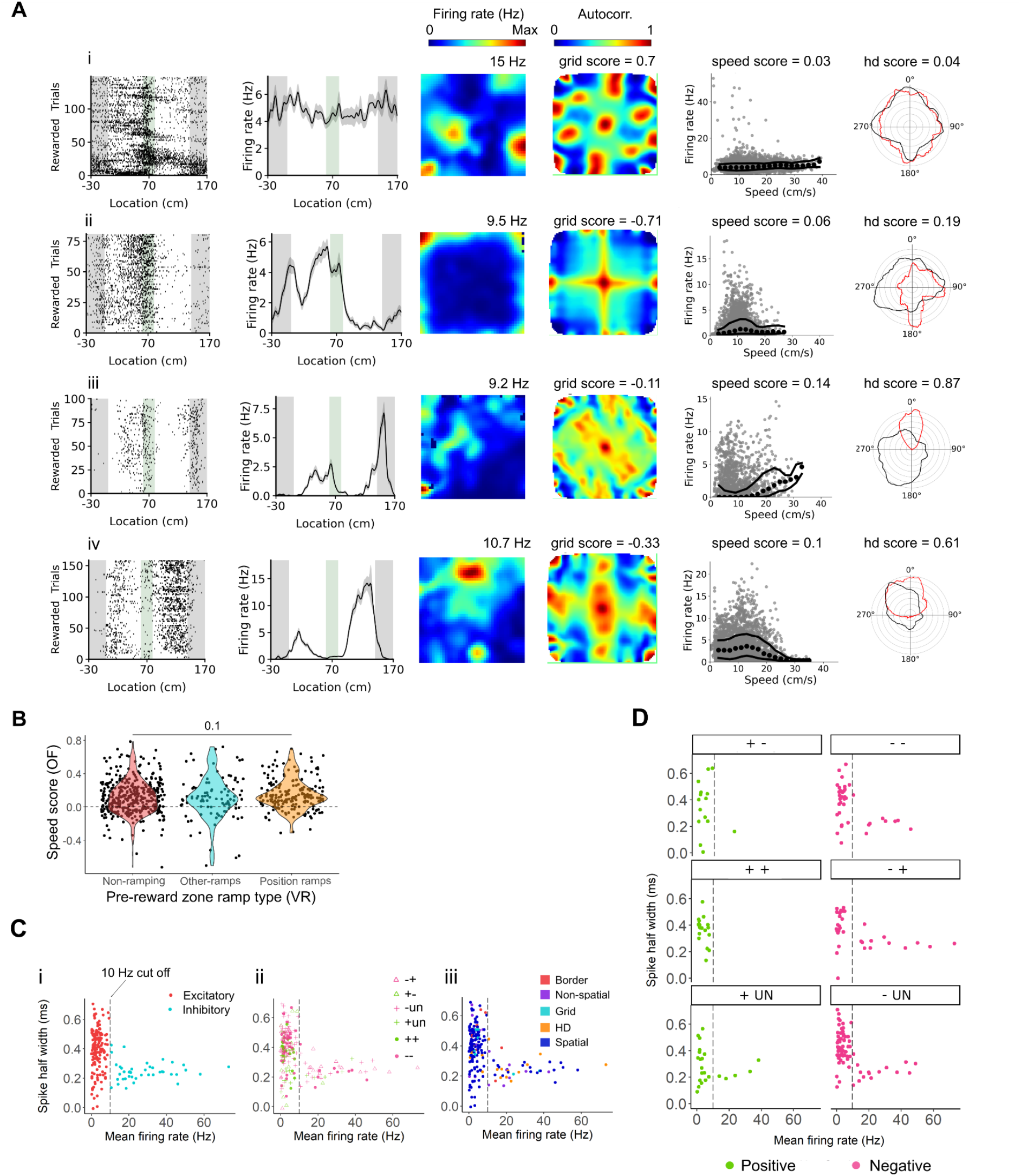
Firing properties of ramp cells in the open arena. (A) Examples of firing profiles in the virtual location estimation task and in the open arena for neurons classified on the basis of their activity in the open arena (see Methods) as a grid cell (i), a head direction cell (ii), a border cell with conjunctive modulation by head direction score (iii) and a pure border cell (iv). The plots from left to right show spike rasters (i) and mean firing rate (ii) as a function of position on the linear track, spatial heat maps (iii) and spatial autocorrelograms of spike rate in the open arena, firing rate as a function of speed in the open arena (vii) and polar plots of average firing rate (red) and movement (black) as a function of head direction in the open arena. (B) The distribution of speed scores was similar among positionally modulated ramping neurons, other ramping neurons and non-ramping neurons (ANOVA, DF = 2, F = 2.23, p = 0.10). (C) Scatter spike half width as a function of firing rate for all position encoding ramp neurons (n = 618), colour coded according to classification of putative interneurons (rate > 10 Hz) and excitatory cells (rate < 10 Hz)(i), to ramp firing rate profile approaching and following the reward zone (ii), and to firing classification in the open arena (iii). (D) Scatter plots of spike half width as a function of firing rate for different firing rate profiles during the virtual location estimation task.

## Supplemental Data

**Supplemental Data 1.**
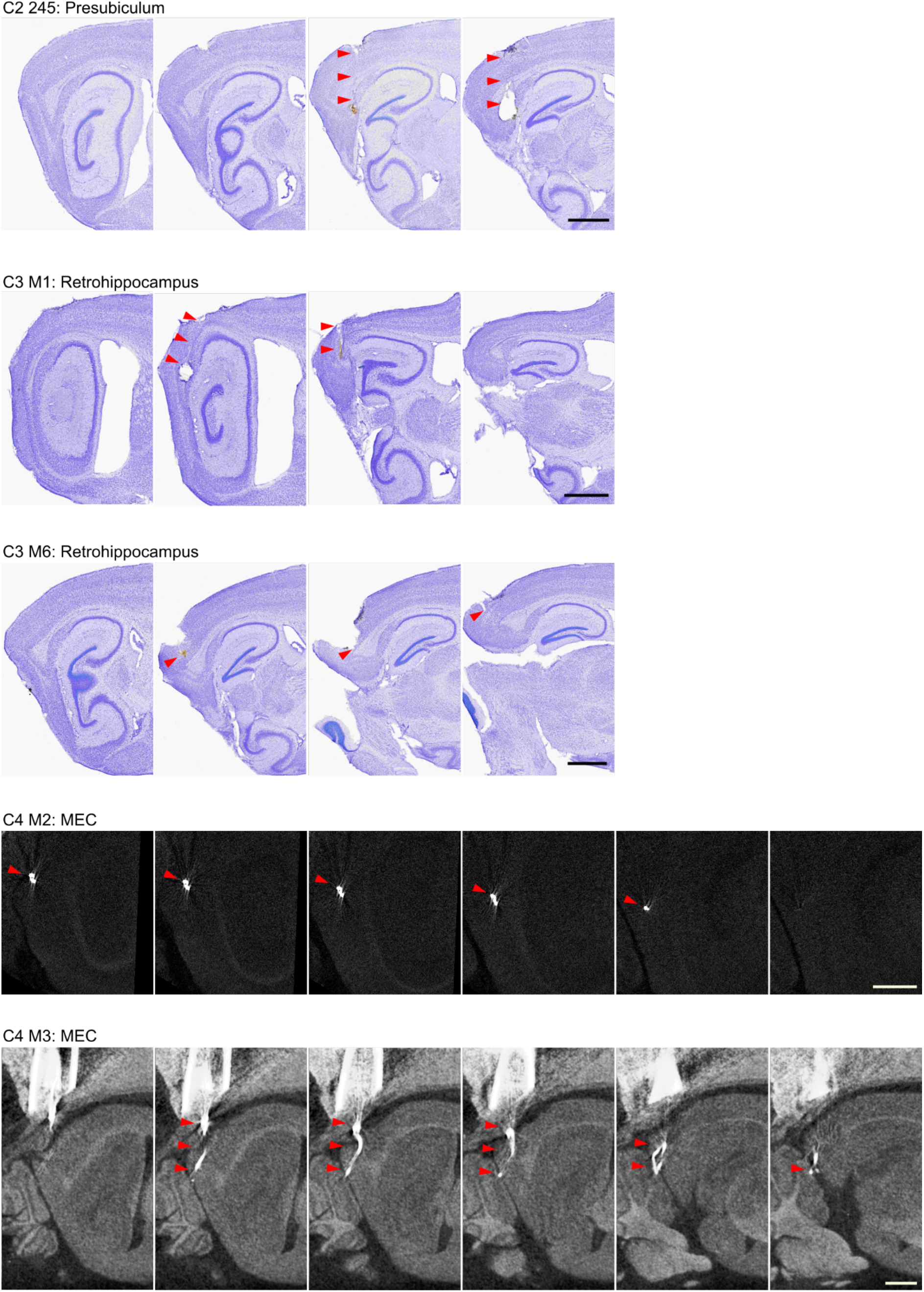
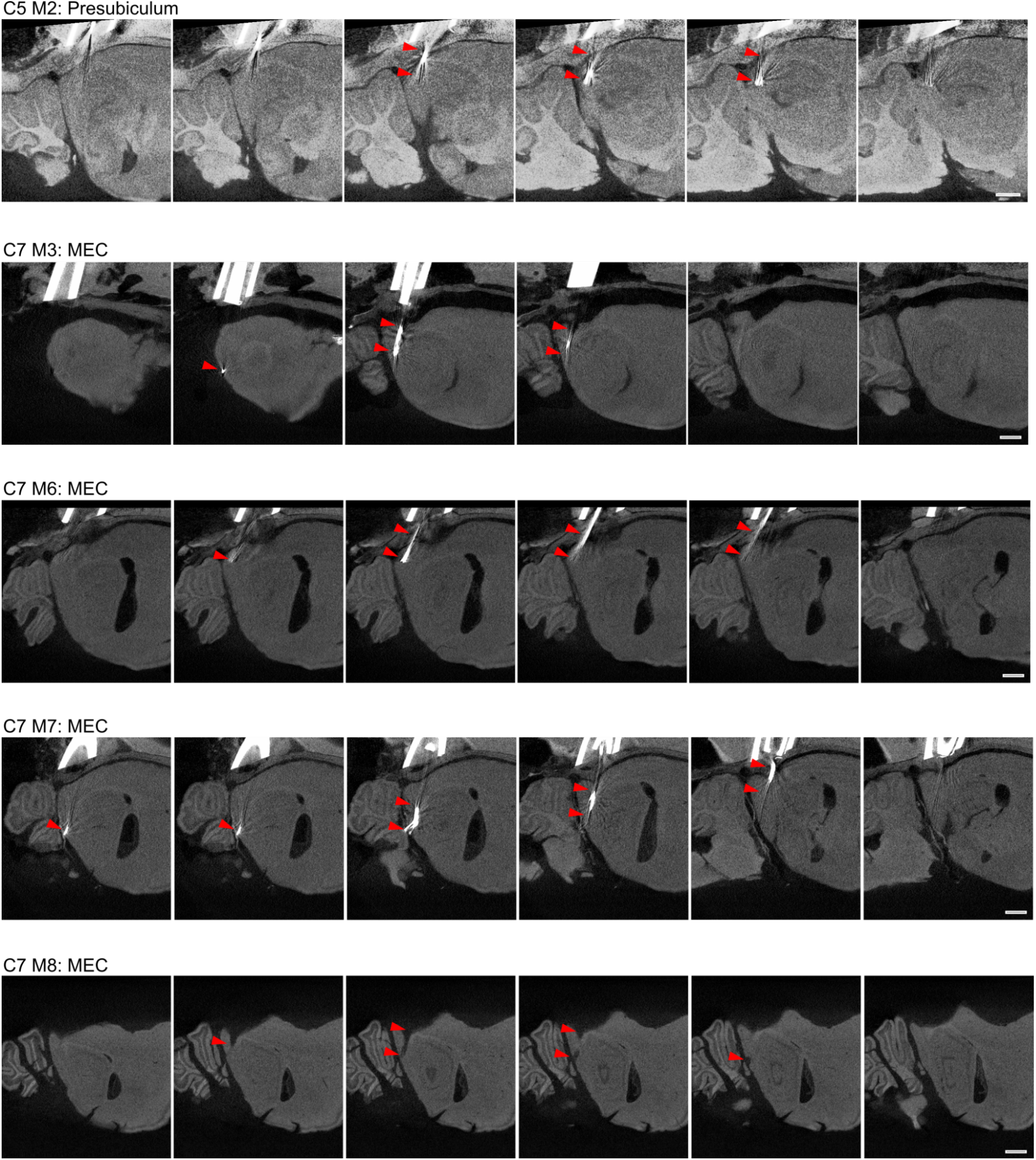
Tetrode localisation. Cresyl violet stained brain sections used for assessment of tetrode locations in three mice and Micro-CT images used for assessment of tetrode locations in six mice. For each animal, sagittal slices are presented lateral to medial from left to right and the classification of the tetrodes target is shown at the top left. This classification is based on the terminal location of the tetrode and the distance travelled during the experiment (see Methods). Two mice had no visible tetrode tracks in any slice and are not shown. In all images red triangles point to the putative tetrode tracks. The number of neurons recorded in each mouse was: C2 245: 47 neurons; C3 M1: 181 neurons; C3 M6: 56 neurons; C4 M2: 314 neurons; C4 M3: 48 neurons; C5 M2: 21 neurons; C7 M3: 266 neurons; C7 M6: 243 neurons; C7 M7: 204 neurons. No histology: C5 M1: 31 neurons; C7 M4: 23 neurons. Scale bar denotes 1 mm.

**Supplemental Table 1:**
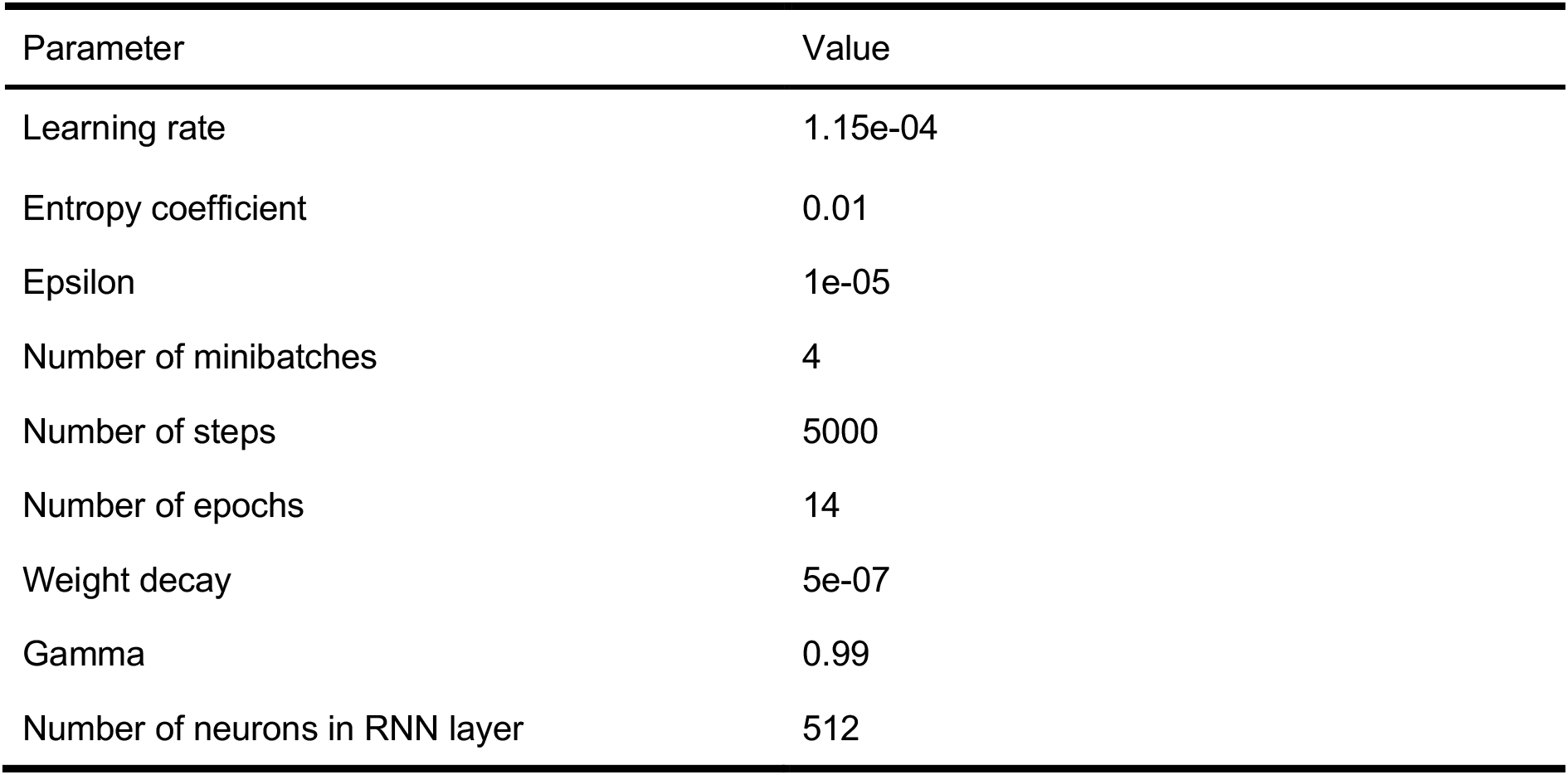
Model hyperparameters. Hyperparameters used in the recurrent neural network model

| Parameter | Value |
| --- | --- |
| Learning rate | 1.15e-04 |
| Entropy coefficient | 0.01 |
| Epsilon | 1e-05 |
| Number of minibatches | 4 |
| Number of steps | 5000 |
| Number of epochs | 14 |
| Weight decay | 5e-07 |
| Gamma | 0.99 |
| Number of neurons in RNN layer | 512 |

